# Human pluripotent stem cell-derived brain pericyte-like cells induce blood-brain barrier properties

**DOI:** 10.1101/387100

**Authors:** Matthew J. Stebbins, Benjamin D. Gastfriend, Scott G. Canfield, Ming-Song Lee, Drew Richards, Madeline G. Faubion, Wan-Ju Li, Richard Daneman, Sean P. Palecek, Eric V. Shusta

**Author notes:** **Correspondence:** Drs. Eric V. Shusta and Sean P. Palecek, Department of Chemical and Biological Engineering, University of Wisconsin-Madison, 1415 Engineering Dr., Madison, WI 53706, United States of America.

## Abstract

Brain pericytes play an important role in the formation and maintenance of the neurovascular unit (NVU), and their dysfunction has been implicated in central nervous system (CNS) disorders. While human pluripotent stem cells (hPSCs) have been used to model other components of the NVU including brain microvascular endothelial cells (BMECs), astrocytes, and neurons, cells having brain pericyte-like phenotypes have not been described. In this study, we generated neural crest stem cells (NCSCs), the embryonic precursor to forebrain pericytes, from human pluripotent stem cells (hPSCs) and subsequently differentiated NCSCs to brain pericyte-like cells. The brain pericyte-like cells expressed marker profiles that closely resembled primary human brain pericytes, and they self-assembled with endothelial cells to support vascular tube formation. Importantly, the brain pericyte-like cells induced blood-brain barrier (BBB) properties in BMECs, including barrier enhancement and reduction of transcytosis. Finally, brain pericyte-like cells were incorporated with iPSC-derived BMECs, astrocytes, and neurons to form an isogenic human NVU model that should prove useful for the study of the BBB in CNS health, disease, and therapy.

## INTRODUCTION

The blood-brain barrier (BBB) is comprised of specialized brain microvascular endothelial cells (BMECs) that line the vasculature of the central nervous system (CNS). BMECs allow for the selective passage of essential nutrients and metabolites into the brain and help prevent the entry of damaging substances. While the BBB plays an important role in CNS homeostasis, it also creates a bottleneck for the delivery of therapeutics (*1–3*). In addition, BBB dysfunction has been observed in many CNS pathologies including Alzheimer’s disease, multiple sclerosis, and stroke, and increasing evidence demonstrates treating BBB contribution to CNS disorders may improve disease outcomes (*4–12*). Importantly, BMECs gain their unique properties as a result of coordinated signaling cues from other brain cells surrounding CNS microvessels, including CNS pericytes, astrocytes, and neurons that together with BMECs form the neurovascular unit (NVU) (*13–18*). Recently, brain pericyte contributions to BBB development and function have begun to be elucidated, and potential pericyte roles in CNS disease have been suggested. CNS pericytes associate with BMECs early in embryonic development as nascent blood vessels invade the developing neural tube. The emergence of pericytes corresponds to BBB formation through reduction of transcytosis, decreased immune cell adhesion molecule expression, and reduced ultrastructural tight junction abnormalities (*13*). In the adult, pericytes regulate vascular stability and diameter (*5, 19–21*), contribute to the BMEC basement membrane (*20, 22–24*), regulate BMEC molecular phenotype (*14, 25*), and reduce transcytosis (*14*).

As a result of the emerging importance of brain pericytes in brain health and disease, they have been increasingly incorporated into *in vitro* models of the BBB. For example, co-culture with pericytes can improve BMEC phenotype in co-culture systems, inhibit endothelial cell tube formation *in vitro* (*26*), and induce BMEC properties in hematopoietic stem cell-derived endothelial cells (*27–29*). We also reported that primary brain pericytes could be combined with human pluripotent stem cell derived BMECs (hPSC-derived BMECs) and enhance their functionality (*30*). Such hPSC-derived BBB models offer the capability for screening of CNS-penetrant therapeutics (*37*) and can be used to investigate BBB contributions to human disease using patient-derived induced pluripotent stem cells (iPSCs) (*32, 33*). While we and others have recently demonstrated the combination of iPSC-derived BMECs with iPSC-derived astrocytes and neurons to form high fidelity multicellular BBB models (*34–36*), the inclusion of pericytes, to date, has been limited to primary human sources (*30, 35*). Unfortunately, primary sources do not scale with high fidelity (*37, 38*), and unlike iPSC sources, do not reflect the genetic contributions that can be important to modeling human disease. Thus, for patient-specific modeling of the healthy and diseased BBB, it is paramount to generate brain pericyte-like cells from human iPSCs.

Vascular mural cells include both smooth muscle cells, which line arterioles and venules, and pericytes, which are associated with smaller microvessels and capillaries. Until very recently, it has been difficult to distinguish smooth muscle cells from pericytes based on marker expression (*39*). Moreover, hPSC-derived mural cells from different embryonic origins display functionally distinct phenotypes and respond differentially to disease pathways (*40, 41*). While most mural cells originate from mesoderm, CNS forebrain mural cells arise from neural crest stem cells (NCSCs) (*42, 43*), a multipotent stem cell population capable of forming peripheral neurons and mesenchymal derivatives including adipocytes, osteocytes, and chondrocytes (*44, 45*), among other cell types. Previous studies have described processes to differentiate hPSCs to NCSCs and demonstrated their potential to form vascular smooth muscle cells (*41, 45, 46*).

However, it is unknown whether NCSCs can generate pericyte-like cells that enhance BBB phenotypes in BMECs. Here, we describe a facile protocol for generating multipotent NCSCs from hPSCs by canonical WNT signaling activation with simultaneous inhibition of BMP and activin/nodal signaling as previously described (*45, 47*). These hPSC-derived NCSCs can be further differentiated to mural cells that express pericyte markers by 9 days of culture in serum-containing medium. These pericyte-like cells associated with vascular tube networks and induced key pericyte-driven phenotypes in BMECs including the enhancement of barrier properties and reduction of transcytosis. Finally, an isogenic model of the NVU comprised of iPSC-derived pericytes, BMECs, astrocytes, and neurons, exhibited elevated barrier properties compared to a model lacking pericytes, suggesting future applications of iPSC-derived pericytes in CNS drug screening, BBB development studies and disease modeling applications.

## RESULTS

### Directed differentiation of hPSCs to NCSCs in low protein medium

We first assessed the capability of E6, a reduced factor medium, to support differentiation of H9 human embryonic stem cells (hESCs) and IMR90C4 and CS03n2 induced pluripotent stem cells (iPSCs) to NCSCs. H9 hESCs were cultured for 15 days in E6 medium supplemented with heparin and pathway modulators previously implicated in hPSC differentiation to NCSCs (*47*): 1 μM CHIR99021, a GSK3β inhibitor to promote WNT signaling; 10 μM SB431543, an ALK5 antagonist to inhibit Activin/Nodal/TGFβ signaling; and 10 ng/mL FGF2 (E6-CSF). However, E6-CSF failed to produce p75-NGFR^+^/HNK1^+^ NCSCs, and increasing CHIR99021 concentration (2 μM) did not aid in inducing p75-NGFR expression (Figure S1A).

BMP signaling during hPSC differentiation to NCSC can inhibit NCSC formation, and WNT signaling activation can induce downstream BMP signaling in hPSCs (*45*); however, the requirement of BMP inhibition in NCSC differentiation has been variable (*41, 45*). To examine the effects of BMP inhibition on hPSC differentiation to NCSCs in minimal medium, E6-CSF medium was supplemented with 1 μM dorsomorphin, a BMP type I receptor inhibitor, to generate E6-CSFD. With BMP inhibition, H9 hESCs progressed to p75-NGFR^+^/HNK1^+^ NCSCs that also expressed AP-2 after 15 days of E6-CSFD treatment (Figure 1A-C, Figure S1A, Table 1). E6-CSFD also drove NCSC formation in IMR90C4 and CS03n2 iPSC lines (Figure S1B-E, Table 1). H9 and CS03n2 hPSCs yielded cultures comprising ~90% NCSCs, while purity of IMR90C4-derived NCSCs was frequently lower (Table 1). Temporal mRNA analysis confirmed loss of pluripotency by D15 of E6-CSFD treatment, as indicated by loss of *NANOG* and *POU5F1* pluripotency transcripts (Figure 1D, Figure S1E). In addition, after 15 days of E6-CSFD treatment, the differentiation mixture expressed NCSC-associated transcripts, including *TFAP2A, SOX9, SOX10, B3GAT1* (HNK1) and *NGFR* (Figure 1D, Figure S1E). At D15 of E6-CSFD treatment, cells had undergone approximately 7 population doublings (Figure 1E), corresponding to over 100 NCSCs per input hPSC.

**Figure 1:**
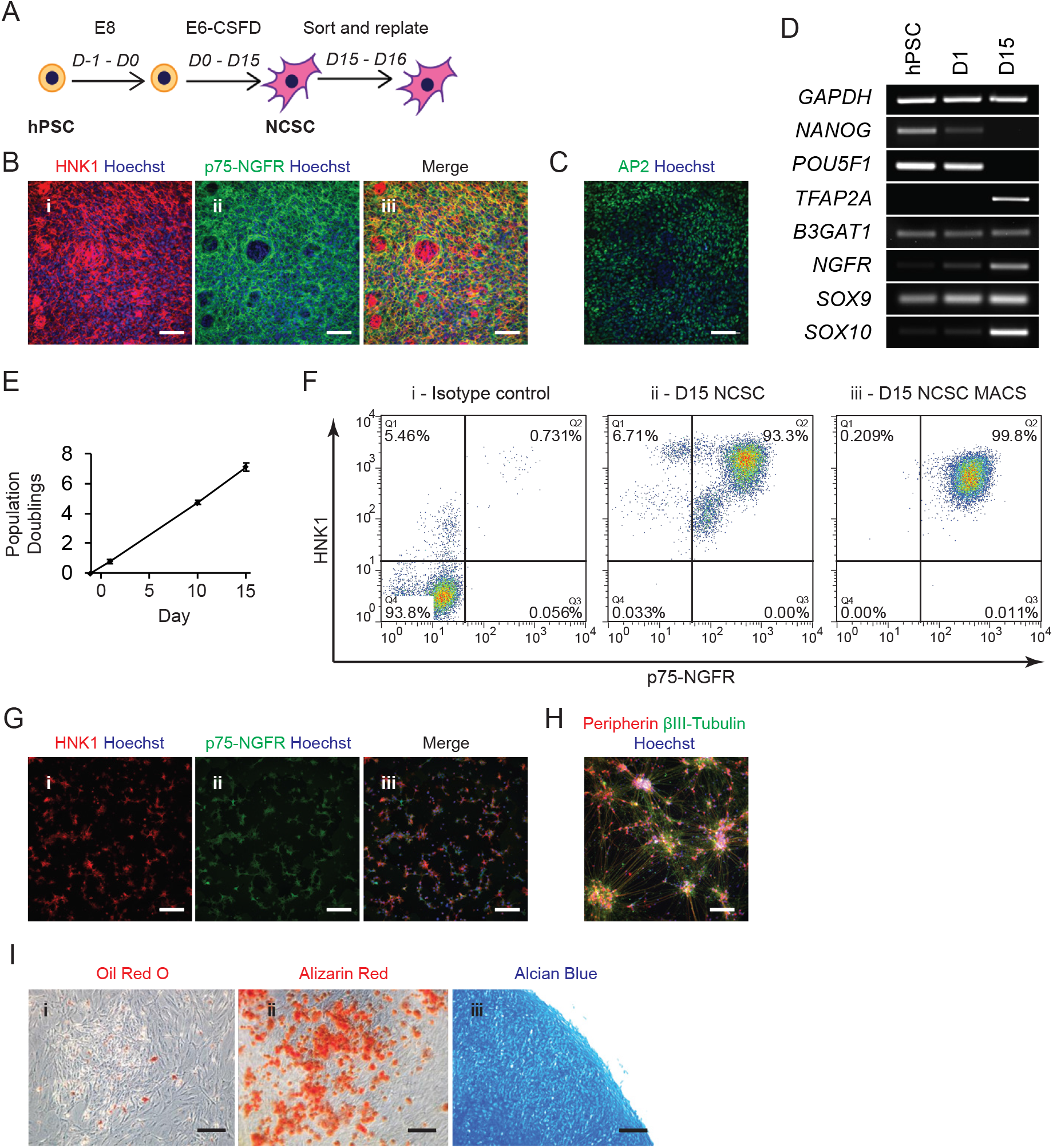
Generation of multipotent NCSC populations. A) NCSC differentiation timeline. Small molecule activation of canonical Wnt signaling and small molecule inhibition of Activin/Nodal/TGFβ/BMP signaling in minimal medium produces H9-derived NCSCs over a 15 day treatment window. NCSCs are then magnetically sorted and replated for subsequent mural cell differentiation. B) Immunocytochemistry images of H9 hESCs differentiated in E6-CSFD probed for the presence of HNK1 and p75-NGFR at D15. NCSCs are HNK1^+^/p75-NGFR^+^ cells. Hoechst nuclear counter stain (blue) is also included. Scale bar: 100 μm. C) AP-2 immunocytochemistry images for H9-derived NCSCs at D15. Hoechst nuclear counter stain (blue) is also included. Scale bar: 100 μm. D) Temporal PCR analysis of pluripotency (*NANOG, POU5F7*) and NCSC (*TFAP2A, B3GAT7, NGFR, SOX9, SOX70*) transcripts. E) Quantification of NCSC expansion in population doublings over the 15 days of NCSC differentiation. Plotted are the mean ± SD of three technical replicates of a representative differentiation. F) Flow cytometry analysis of H9-derived NCSCs. Panels include isotype controls (panel i), NCSC (HNK1+/p75-NGFR+) purity prior to MACS (panel ii), and NCSC purity following MACS (panel iii). Inset percentages are included in each quadrant. Quantitation is shown in Table 1. G) Immunocytochemistry analysis of D16 NCSCs following MACS and replating. NCSCs maintained HNK1 and p75-NGFR expression. Hoechst nuclear counter stain (blue) is also included. Scale bar: 100 μm. H) Immunocytochemistry analysis of H9-derived NCSCs subsequently differentiated in peripheral neuron medium. Resultant cells were positive for βIII-tubulin and peripherin expression. Hoechst nuclear counter stain (blue) is also included. Scale bar: 200 μm. I) H9-derived NCSCs could be differentiated into mesenchymal derivatives, including Oil Red O stained adipocytes (panel i, red), Alizarin red stained osteocytes (panel ii, red), and Alcian blue stained chondrocytes (panel ii, blue). Scale bar: 200 μm.

**Table 1:**
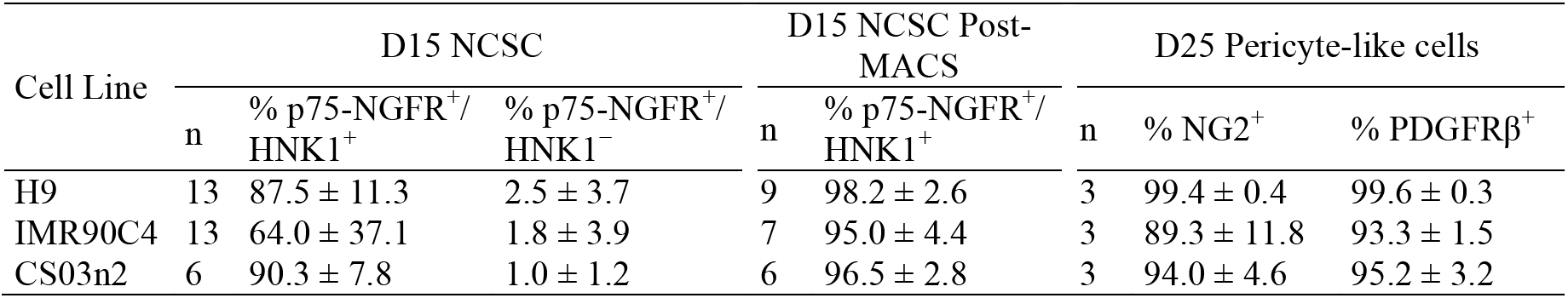
NCSC and pericyte-like cell differentiation efficiency.

To purify NCSCs from the differentiation cultures, day 15 NCSCs were positively selected using anti-p75-NGFR magnetic activated cell sorting (MACS). MACS enriched p75-NGFR^+^/HNK1^+^ NCSC populations above 95% for all three hPSC lines tested (Figure 1F, Figure S2A-B, Table 1). Sorted NCSCs retained p75-NGFR and HNK1 expression following replating (Figure 1G). In addition, treating NCSCs with N2 medium supplemented with BDNF, GDNF, NT-3, and NGF-β yielded βIII-tubulin+/peripherin+ peripheral neurons (Figure 1H, Figure S2D). We additionally expanded sorted NCSCs for 11 days and then differentiated these cells to mesenchymal derivatives: Oil Red O+ adipocytes were obtained by treating NCSCs with insulin, IBMX, and dexamethasone, Alcian blue+ chondrocytes using pellet culture and TGFβ1-containing chondrogenic medium, and Alizarin red+ osteocytes using dexamethasone, glycerophosphate, and ascorbic acid (Figure 1I; Figure S2C). Taken together, these data demonstrate that reduced factor, low protein E6-CSFD medium directs hPSCs to NCSCs over a 15-day differentiation period, and that MACS-purified NCSCs retain the potential to form NCSC derivatives.

### Serum treatment directs hPSC-derived NCSCs to mural cell lineages

We subsequently identified differentiation conditions capable of driving NCSCs to mural cell lineages (Figure 2A), as defined by coexpression of PDGFRβ and NG2 (*39, 48*). PDGFRβ was expressed in D15 NCSCs (Figure 2C) and in replated cells one day following MACS (D16), but NG2 expression was absent in both of these cell populations (Figure 2C,F). Given the importance of PDGF-BB and TGFβ1 in mural cell development (*49, 50*), we first tested if these factors could induce NG2 expression in NCSCs while also maintaining PDGFRβ expression. Culture of NCSCs for six days in E6 medium generated cells that were PDGFRβ positive but NG2 expression was not observed (Figure 2B). Supplementation of E6 medium with PDGF-BB and TGFβ1 did not induce NG2 expression. However, when E6 medium was supplemented with 10% FBS, resultant cells expressed both PDGFRβ and NG2 (Figure 2B). Comparing differentiation in E6 + 10% FBS on uncoated tissue culture polystyrene (TCPS) to gelatin-coated TCPS, which has previously been reported as conducive to mural cell differentiation (*51*), the uncoated substrate yielded a qualitatively larger fraction of cells that expressed PDGFRβ and NG2 (Figure 2B). Given the capacity for E6 + 10% FBS on uncoated substrate to direct hPSC-derived NCSCs to PDGFRβ+/NG2+ mural cells, we further evaluated these cells.

**Figure 2:**
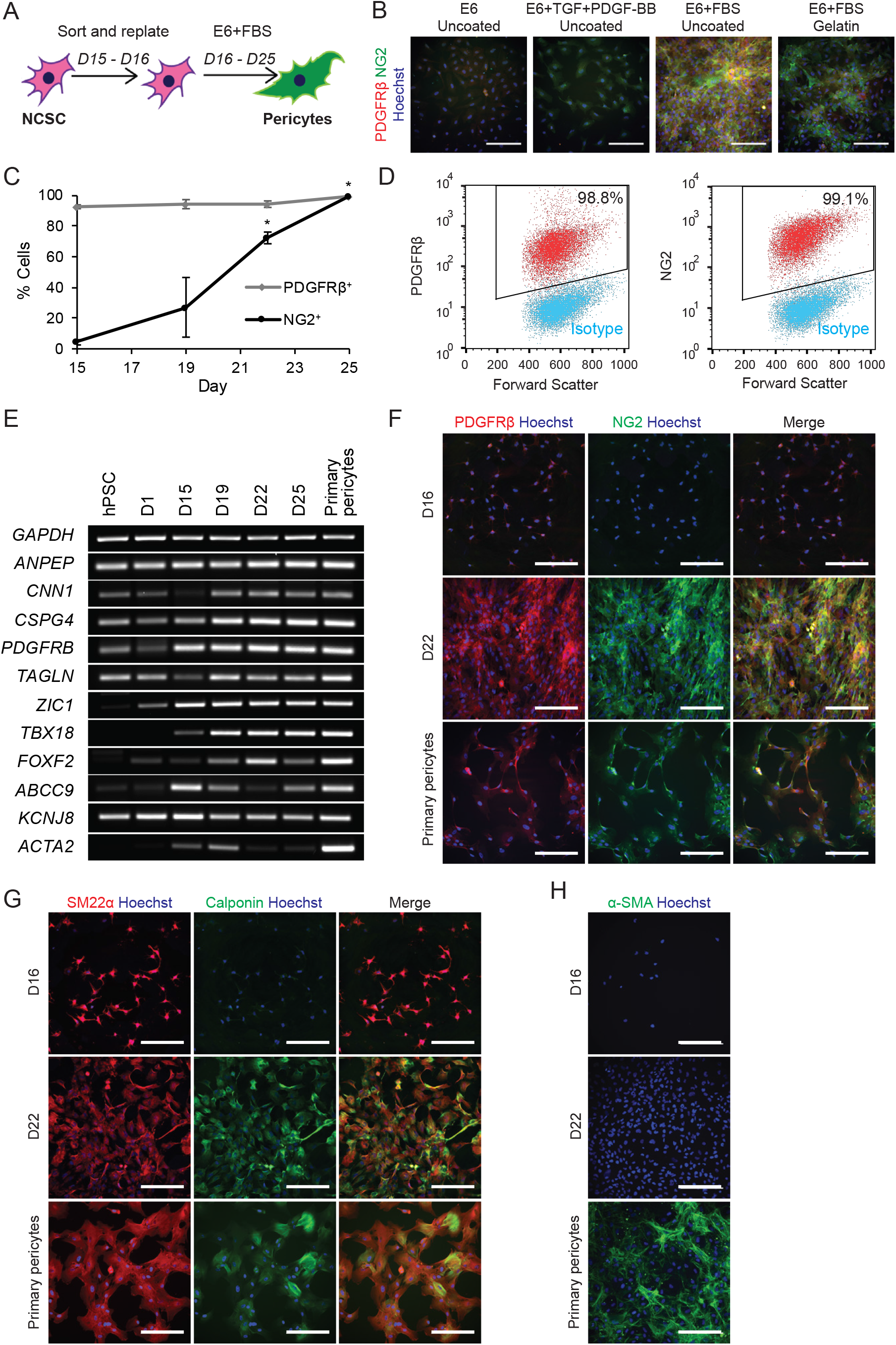
Serum treatment directs H9-derived NCSCs towards mural cells. A) Differentiation timeline for mural cell differentiation. Replated NCSCs are differentiated to mural cells in E6 medium plus 10% FBS for 9 days. B) PDGFRβ and NG2 immunocytochemistry of cells obtained after treating replated H9-derived NCSCs for 6 days in E6, E6 + TGFβ1 + PDGF-BB, or E6 + 10% FBS on uncoated tissue culture polystyrene, or E6 + 10% FBS on gelatin-coated tissue culture polystyrene. C) Temporal flow cytometry analysis for PDGFRβ and NG2 positive cells in H9-derived NCSCs treated with E6 + 10% FBS. Depicted are the means ± SEM of at least two independent differentiations at each time point, * *P* < 0.05 vs. D15 NCSC using ANOVA followed by Dunnett’s test. D) Representative PDGFRβ and NG2 flow cytometry plots for H9-derived NCSC treated 9 days with E6 + 10% FBS medium. Quantitative data can be found in Table 1. E) Temporal PCR analysis of mural and pericyte transcripts for the differentiating H9 hESCs. F) PDGFRβ and NG2 immunocytochemistry of H9-derived NCSCs (D16), mural cells (D22), and primary pericytes. Hoechst nuclear counter stain (blue) is also included. Scale bar: 200 μm. G) Calponin and SM22α immunocytochemistry of H9-derived NCSCs (D16), mural cells (D22) and primary pericytes. Hoechst nuclear counter stain (blue) is also included. Scale bar: 200 μm. H) α-SMA immunocytochemistry of H9-derived NCSCs (D16), mural cells (D22) and primary pericytes. Hoechst nuclear counter stain (blue) is also included. Scale bar: 200 μm.

The temporal evolution of hPSC-derived NCSCs to PDGFRβ+/NG2+ mural cells using E6 + 10% FBS was examined over a 9 day period (D16-D25). At D15 of differentiation, 92.4 ± 1.1% of H9-derived NCSCs expressed PDGFRβ; and after 9 days of serum treatment, nearly all cells were PDGFRβ+ (99.6 ± 0.2%) (Figure 2C-D), with expression of *PDGFRB* transcript present in D15 NCSCs and throughout the differentiation in serum (Figure 2E). In contrast, despite the fact that the NG2-encoding *CSPG4* transcript was expressed in D15 NCSCs (Figure 2E), NG2 protein was not detected at this time point by flow cytometry (Fig 2C). However, the percentage of cells expressing NG2 increased over the 9 day differentiation period, with nearly all cells becoming NG2+ (99.4 ± 0.3% at D25, *P* < 0.05 vs. D15) (Figure 2C-D). The E6 + 10% FBS differentiation scheme also generated at least ~90% PDGFRβ+ and NG2+ cells in IMR90C4- and CS03n2-derived NCSCs following nine days of E6 + 10% FBS treatment (D25; Table 1; Figure S3A-D). At D22, this procedure yielded a roughly ten-fold expansion in mural cells (9.5 ± 1.3 mural cells per sorted NCSC for six independent differentiations).

To further probe the transition of hPSC-derived NCSCs to pericyte-like cells, we examined the temporal evolution of transcripts that have been associated with pericytes and other mural cells. H9 hESCs expressed *CNN1* (calponin) and *TAGLN* (SM22α), which encode contractile proteins implicated in early mural cell differentiation (*40*). Upon NCSC differentiation, transcript abundance was reduced (D15), while subsequent serum treatment elevated expression of these transcripts (Figure 2E). At D16, differentiating hPSC-derived NCSC expressed SM22a but calponin expression was not observed. (Figure 2G, Figure S3F,I). By D22, differentiating hPSC-derived NCSCs exhibited calponin/SM22α coexpression with cellular localization to contractile fibers (Figure 2G, Figure S3F,I). Interestingly, smooth muscle actin (α-SMA) was not detected in D22 cells treated with E6 + 10% FBS, although serum transiently increased abundance of the transcript (*ACTA2*) before downregulation (Figure 2E,H; Figure S3G,J; Figure S6). In contrast, NCSCs treated with E6 alone or E6 plus PDGF-BB and TGFβ1 expressed α-SMA in addition to calponin and SM22α (Figure S4). In addition, these cells exhibited a morphology similar to smooth muscle cells, with large cell bodies and distinct cell borders, whereas the cells differentiated in E6 + 10% FBS were smaller with numerous projections reminiscent of cultured primary brain pericytes (Figure 2G-H). After extended culture in E6 + 10% FBS (D45), the resultant cells continued to be PDGFRβ+/NG2+ and expressed calponin and SM22α while α-SMA was still absent (Figure S5). Primary human brain pericytes expressed all three contractile proteins, and had a morphology similar to the serum treated NCSC-derived mural cells (Figure 2F-H).

Additional transcript analysis was used to further characterize the differentiation process. The mural cell marker, *ANPEP* (CD13), was expressed throughout the differentiation process. While *PDGFRB, CSPG4* (NG2), *CNN1, TAGLN, ANPEP*, and *TBX18* are mural cell markers expressed throughout the body, *FOXF2* and *ZIC1* have been suggested as being selectively expressed in brain mural cells (*52–54*). Accordingly, given the NCSC origin of the mural cells *FOXF2* and *ZIC1* were induced during the differentiation although *FOXF2* expression decreased at D25 in H9-derived cells (Figure 2E, Figure S6). Until recently, it has been difficult to use markers to distinguish pericytes from smooth muscle cells in brain; however, it has been suggested that *ABCC9* and *KCNJ8* are two transcripts having selective expression in brain pericytes as compared to smooth muscle (*39, 48*). *ABCC9* levels were biphasic with strong expression in D15 NCSC and then a re-induction in D25 mural cells. *KCNJ8* was expressed fairly uniformly throughout the differentiation process (Figure 2E). Similar results were observed for mural cells derived from IMR90C4- and CS03n2-derived NCSCs, although the IMR90C4 mural cells had weaker *ZIC1* and *ABCC9* signatures (Figure S6). Overall, the transcript profile of mural and pericyte-associated genes in the NCSC-derived mural cells was qualitatively very similar to that of primary human brain pericytes (Fig 2E).

We next used RNA-sequencing (RNA-seq) to quantify global gene expression in NCSC-derived mural cells and to evaluate the temporal emergence of a pericyte-like population. As expected, unbiased hierarchical clustering based on expression (fragments per kilobase of transcript per million mapped reads, FPKM) of all transcripts revealed the highest similarity between NCSC-derived mural cells generated from three independent differentiations from H9 hESCs as well as the two differentiations from IMR90C4 and CS03n2 iPSCs (Figure 3A, D25 sample cluster). The Pearson correlation coefficients comparing transcript expression in H9-derived mural cells at D25 to the two replicate H9 differentiations were 0.99 and 0.98 (D25 H9-A versus D25 H9-B or H9-C, *P* < 0.0001). Moreover, the Pearson correlation coefficients comparing the mural cells derived from the H9 hESC line to those derived from IMR90C4 and CS03n2 iPSCs were both 0.97 (D25 H9-A versus D25 IMR90 or CS03, *P* < 0.0001). Collectively, these data indicate a highly reproducible differentiation procedure amongst replicated differentiations and hPSC lines. Furthermore, NCSC-derived mural cells at D25 clustered more closely with primary brain pericytes than with NCSCs or hPSCs (Figure 3A). The Pearson correlation coefficient between the average transcript expression of all D25 NCSC-derived mural cell samples and the average of the primary pericyte samples was 0.89 (*P* < 0.0001), suggesting strong positive association between NCSC-derived mural cells and primary human pericytes. Consistent with RT-PCR experiments (Figure 1D, Figure 2E), temporal analysis of transcript expression demonstrated downregulation of pluripotency markers *NANOG* and *POU5F1*, and transient upregulation of *NGFR, B3GAT1, SOX9*, and *S0X10* in D15 NCSCs. We also observed gradual induction of *CSPG4, PDGFRB, CNN1, TAGLN, ANPEP, TBX18, ABCC9*, and *KCNJ8* over the time course of E6 + 10% FBS treatment, and transient upregulation of *ACTA2* and *FOXF2* (Figure 3B). Expression levels of *CSPG4*, *PDGFRB*, CNN1, *TAGLN*, *FOXF2, ABCC9*, and *KCNJ8* were similar in NCSC-derived mural cells and primary brain pericytes; however, consistent with the lack of α-SMA expression (Figure 2H, Figure S3), NCSC-derived mural cells expressed nearly 100-fold less *ACTA2* transcript than primary pericytes (Figure 3B). By D45, NCSC-derived mural cells retained expression of most markers at levels similar to D25 cells, while *ANPEP, ABCC9*, and *KCNJ8* expression further increased, suggesting these cells may continue to mature during extended culture in E6 + 10% FBS (Figure 3B). Comparison of transcripts upregulated in NCSC-derived mural cells compared to their NCSC precursors revealed several enriched Gene Ontology (GO) terms including vascular development, blood vessel morphogenesis, and extracellular matrix organization (Figure 3C), indicating that differentiation is driving the progression from NCSCs to mural cells with vascular-associated transcript signatures. Of the 46 genes with human homologs identified as pericyte-enriched by single cell RNA-seq in mice (*39*), 29 were expressed at or above 1 FPKM by NCSC-derived mural cells and 26 by primary pericytes (Figure 3D, Table S3). Collectively, these data demonstrate that differentiation of NCSCs in E6 + 10% FBS yielded a mural cell population that expressed pericyte-associated markers while closely mimicking primary brain pericytes at a transcriptome level. Thus, we refer to the NCSC-derived mural cells as brain pericyte-like cells throughout the remainder of the manuscript.

**Figure 3:**
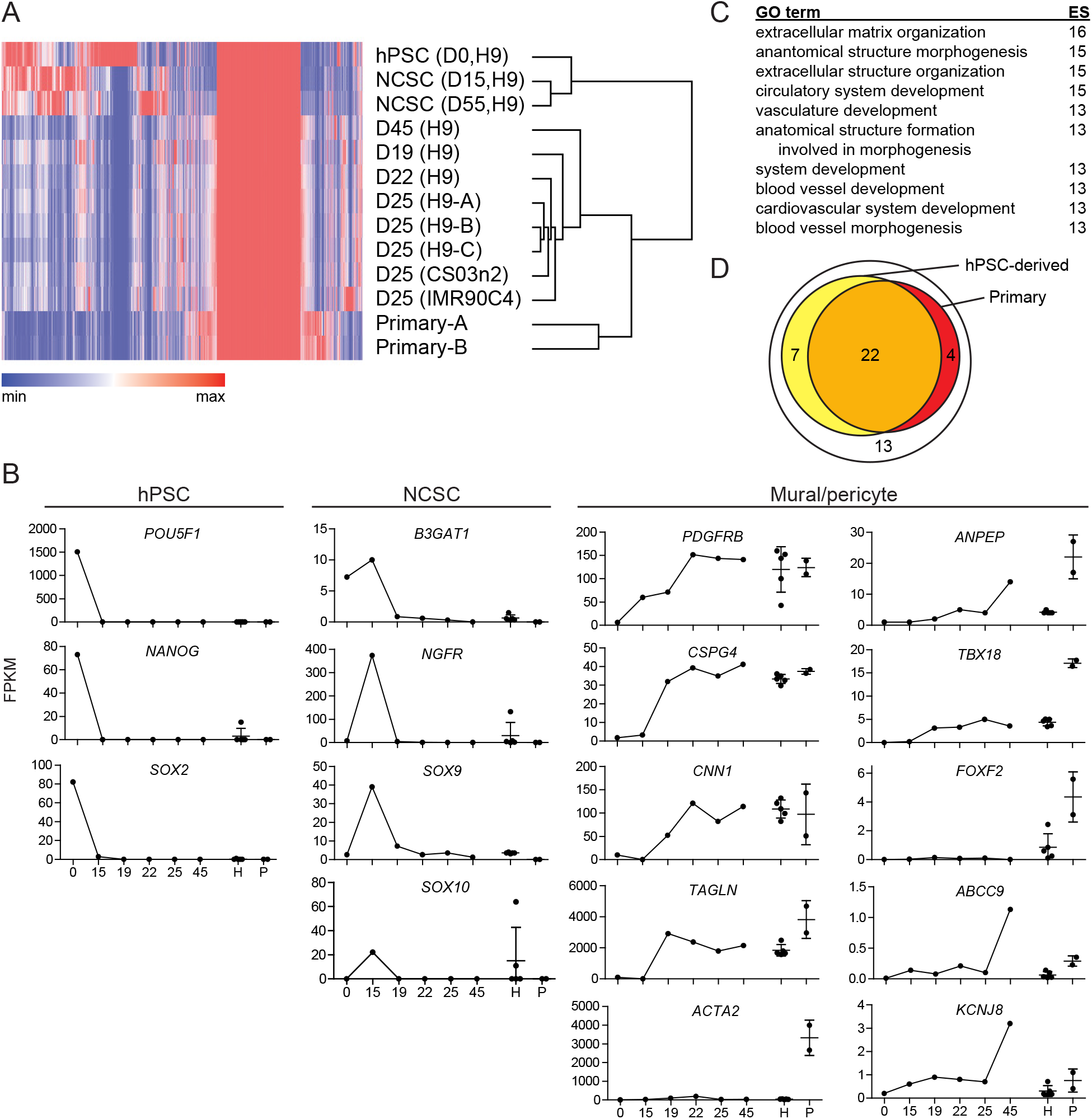
RNA-sequencing of pericyte-like cells and related cell types. A) Hierarchical clustering based on all transcripts of undifferentiated H9 hESCs, H9-derived NCSCs at D15 and after an additional 40 days in E6-CSFD (D55), H9-derived pericyte-like cells at D19, D22, and D25 (three independent differentiations at the D25 time point, indicated as “H9-A”, “H9-B”, and “H9-C”), H9-derived pericyte-like cells maintained for an additional 20 days in E6 + 10% FBS (D45), CS03n2- and IMR90C4-derived pericyte-like cells at D25, and primary brain pericytes (from two distinct cultures of the same cell source, indicated as “Primary-A” and “Primary-B”). B) Expression (FPKM) of selected transcripts in H9 hPSCs (day “0”), NCSCs (“15”), and during the differentiation of pericyte-like cells (“19”, “22”, “25”, and “45”). Also shown is the mean transcript expression in all D25 hPSC-derived pericyte-like cells (H9 A-C, CS03n2 and IMR90C4, “H”) and in primary brain pericytes (“P”). Error bars represent SEM of five independent differentiations (“H”) or of two primary pericyte samples (“P”). C) Top 10 gene ontology (GO) terms, sorted by enrichment score (ES = -log_10_(FDR)), for hPSC-derived pericyte-like cells. Genes included in the dataset were enriched in pericyte-like cells (average of all D25 samples) compared to NCSCs (average of D15 and D55 samples) (FPKM_pericyte-likecells_/FPKM_NCSC_ ≥ 10), and were expressed at ≥ 1 FPKM in pericyte-like cells. D) Expression (≥ 1 FPKM) of murine pericyte-enriched transcripts (46 transcripts (*39*)) in hPSC-derived pericyte-like cells (29 transcripts) and primary brain pericytes (26 transcripts). A detailed listing of genes and FPKM values can be found in Table S3.

### Brain pericyte-like cells assemble with vascular tube networks

Pericytes associate with endothelial cells and stabilize nascent vascular networks (*50*). To assess the ability of brain pericyte-like cells to self-assemble with endothelial cells, an *in vitro* endothelial tube forming assay was performed. A 3:1 mixture of primary pericytes or hPSC-derived brain pericyte-like cells (D22) and human umbilical vein endothelial cells (HUVECs) was plated on Matrigel (Figure 4A). H9, CS03n2 and IMR90C4-derived brain pericyte-like cells self-associated with HUVECs much like primary human brain pericytes (Figure 4B-C). After 24 hours, hPSC-derived pericyte-like cells exhibited high NG2 expression and aligned along the CD31+ endothelial cell tube perimeter and developed pericyte-like morphology with stellate-shaped bodies and extended cell processes (Figure 3B-C). Whereas HUVECs alone and HUVECs in co-culture with control HEK293 cells yielded many small branching tubes, co-culture with the hPSC-derived brain pericyte-like cells or primary human brain pericytes yielded fewer, appreciably longer tubes (Figure 4C-F). These data demonstrate that hPSC-derived NCSC lineage mural cells exhibit pericyte-like association with endothelial cells leading to the formation of more well developed tube networks.

**Figure 4:**
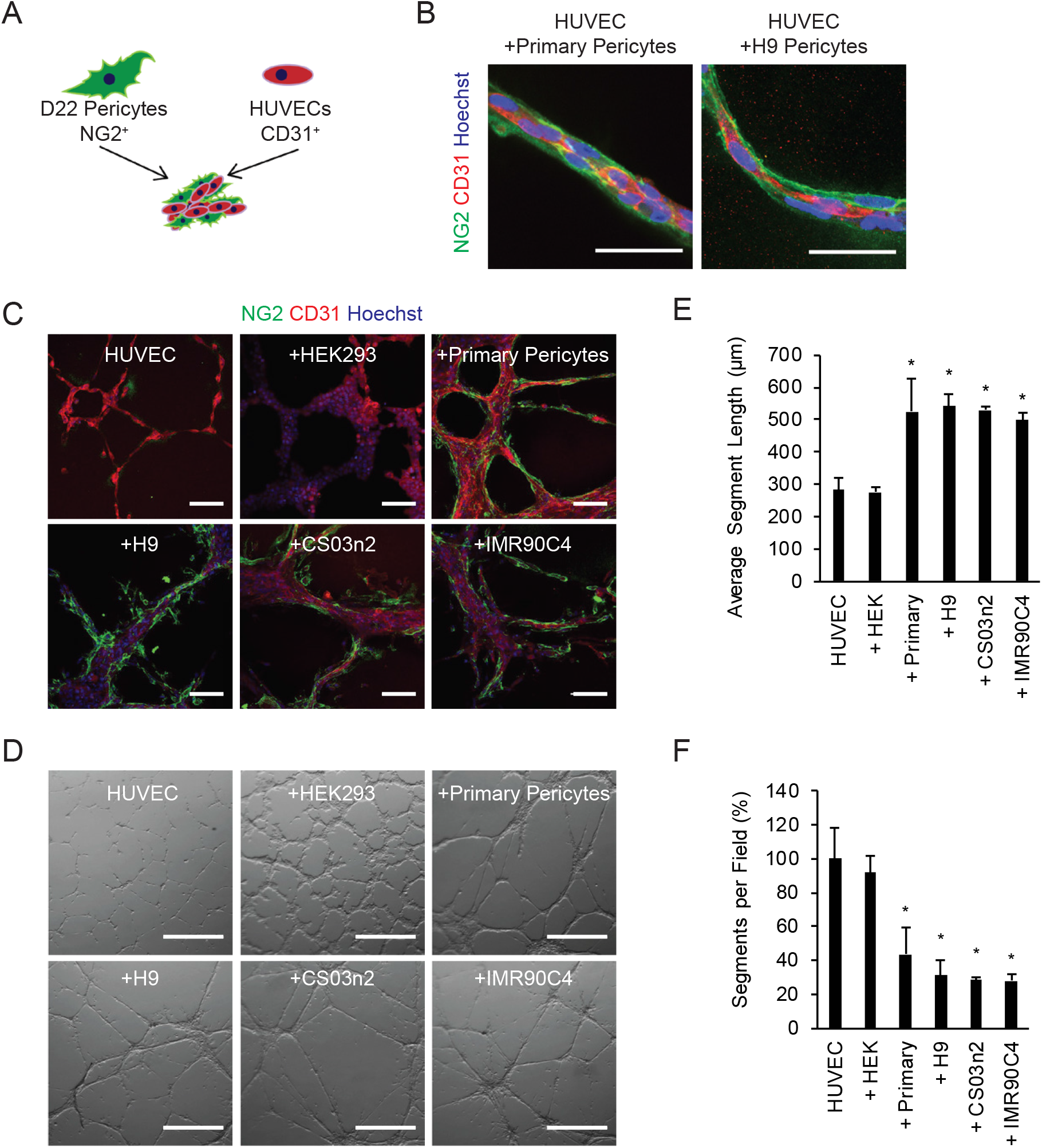
hPSC-derived pericyte-like cell assembly with endothelial cells. A) Self-assembly schematic. hPSC-derived pericyte-like cells self-assemble with HUVECs to form vascular tubes. B) Confocal immunocytochemistry images of primary pericytes and H9-derived pericyte-like cells (NG2) aligning with and extending processes along HUVEC tubes (CD31). Hoechst nuclear counter stain (blue) is also included. Scale bars: 50 μm. C) Immunocytochemistry images of HUVECs alone or cultured with HEK293 fibroblasts (+HEK293), primary human brain pericytes (+Primary Pericytes), CS03n2-derived pericyte-like cells (+CS03n2), H9-derived pericyte-like cells (+H9), or IMR90C4-derived pericyte-like cells (+IMR90C4). Hoechst nuclear counter stain (blue) is also included. Scale bars: 100 μm. D) Representative bright field images of HUVECs alone or cultured with the various cell types. Scale bars: 300 μm. E) Quantification of the average segment lengths from bright field images in panel D. Plotted are means ± SEM of three independent pericyte-like cell differentiations. * *P* < 0.05 vs. HUVEC monoculture; ANOVA followed by Dunnett’s test. F) Quantification of the number of segments per field normalized to HUVEC monoculture from bright field images in panel D. Plotted are means ± SEM of three independent pericyte-like cell differentiations. * *P* < 0.05 vs. HUVEC monoculture; ANOVA followed by Dunnett’s test.

### Brain pericyte-like cells induce blood-brain barrier properties

To investigate if hPSC-derived brain pericyte-like cells can recapitulate key BBB inducing properties that have been observed *in vivo*, such as reduction in tight junction abnormalities and transcytosis, we next co-cultured the pericyte-like cells with hPSC-derived BMECs generated as we previously described (*55*). When D22 brain pericyte-like cells were co-cultured with hPSC-derived BMECs, the BMEC barrier properties as measured by TEER were substantially elevated, while co-culture with a non-inducing cell type (3T3) yielded no barrier enhancement (Figure 5A-B). TEER elevation by hPSC-derived brain-pericyte like cells was indistinguishable from that induced by primary human brain pericytes (Figure 5B). The TEER increases were accompanied by a corresponding decrease in permeability to fluorescein, a hydrophilic, small molecule tracer (Figure 5C). After BMEC co-culture, the brain pericyte-like cells remained NG2+/PDGFRβ+ (Figure S7A), indicating their continued maintenance of mural identity. To determine tight junction changes that may drive the induction in BMEC barrier properties, the expression level and localization of tight junction proteins occludin and claudin-5 were evaluated in the BMECs. Expression levels of occludin and claudin-5 were unchanged by co-culture (Figure S7B). In addition, quantitative immunocytochemical evaluation indicated that the number of cells possessing continuous tight junctions was unchanged upon treatment with pericyte-conditioned medium (Figure 5D-E). However, the percentage of cells with tight junction abnormalities or fraying was substantially reduced by treatment with pericyte-conditioned medium (Figure 4F), correlating with the reduced permeability.

**Figure 5:**
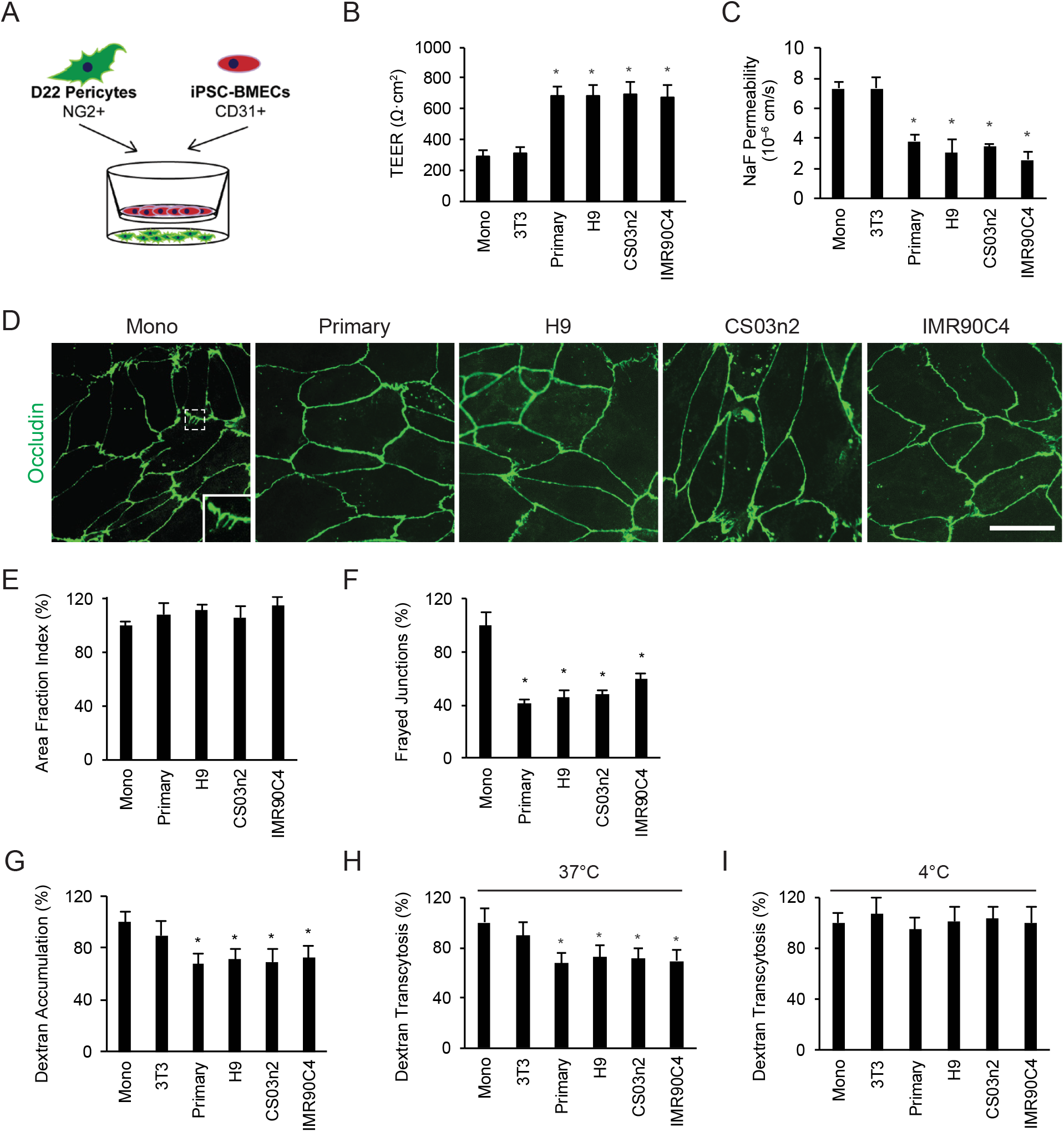
Measurement of the effects of hPSC-derived pericyte-like cells on BBB phenotypes. A) Schematic of Transwell setup for co-culture assays. B) Maximum TEER achieved by IMR90C4-derived BMEC monoculture or co-culture with 3T3 mouse fibroblasts, primary human brain pericytes, H9-derived pericyte-like cells, CS03n2-derived pericyte-like cells, or IMR90C4-derived pericyte-like cells. Plotted are the means ± SEM of at least 3 independent differentiations per condition. * *P* < 0.05 vs. monoculture; ANOVA followed by Dunnett’s test. C) Sodium fluorescein permeability for IMR90C4-derived BMECs in monoculture or co-culture with cell types as described in B. Plotted are the means ± SEM of at least 3 independent differentiations per condition, * *P* < 0.05 vs. monoculture; ANOVA followed by Dunnett’s test. D) Representative images of occludin immunocytochemistry of BMECs cultured for 48 h in EC medium (Mono) or EC medium conditioned by the cell types described in B. Enlarged example of a frayed junction is inset in the monoculture panel. Scale bar: 25 μm. E) Quantification of occludin area fraction index for the samples described in D. Plotted are the means ± SEM of 3 independent differentiations. No significant difference by ANOVA. F) Quantification of frayed junctions visualized by occludin immunocytochemistry for the samples described in D. Plotted are the means ± SEM of 3 independent differentiations. * *P* < 0.05 vs. monoculture; ANOVA followed by Dunnett’s test. G) Accumulation of Alexa-488-tagged 10 kDa dextran in IMR90C4-derived BMECs following 48 hours of co-culture with cell types as described in B. All results are normalized to BMEC monoculture control. Plotted are the means ± SD of 3 Transwells. Results are representative of 3 independent differentiations. * *P* < 0.05 vs. monoculture; ANOVA followed by Dunnett’s test. H,I) Transcytosis of Alexa-488-tagged 10 kDa dextran at 37°C (H) or 4°C (I) across IMR90C4-derived BMECs following 48 hours co-culture with the cell types as described in B. All results are normalized to BMEC monoculture control. Plotted are the means ± SD from 3 Transwells. Results are representative of 3 independent differentiations. * *P* <0.05 vs. monoculture; ANOVA followed by Dunnett’s test. No significant differences at 4°C by ANOVA.

Next, the effects of brain pericyte-like cell co-culture on BMEC transcytosis properties were evaluated. To test non-specific molecular uptake and transcytosis in BMECs, a 10 kDa Alexa 488-tagged dextran was dosed into the apical Transwell chamber and accumulation into and transcytosis across the BMEC monolayer were quantified. After BMEC culture with medium conditioned by hPSC-derived brain pericyte-like cells, confocal imaging indicated a qualitative decrease in intracellular dextran uptake in punctate vesicular structures, similar to that observed with primary human brain pericytes; whereas, medium conditioned by 3T3 control cells had no effect (Figure S7C). Indeed, quantification of dextran accumulation in BMECs co-cultured with brain pericyte-like cells or primary brain pericytes indicated that BMEC accumulation was reduced by about 30% (Figure 5G). These differences in accumulation translated to a corresponding 30% decrease in 10 kDa dextran transcytosis upon pericyte co-culture (Figure 5H). In contrast, when 10 kDa dextran transport was measured at 4°C, conditions that significantly inhibit vesicular transcytosis processes, pericyte co-culture did not affect dextran transport compared to 3T3s or hPSC-derived BMEC monoculture (Figure 5I), indicating that the observed decreases in dextran transport could not be ascribed to differences in paracellular transport resulting from improved tight junction fidelity.

Finally, to confirm that the effects of hPSC-derived brain pericyte-like cells are not specific to BMECs derived from hPSCs, the induction of BBB barrier and transcytosis attributes was also evaluated in primary rat BMECs. Co-culture with IMR90C4-derived brain pericyte-like cells elevated the TEER in primary rat BMECs to the same level as observed with primary human brain pericytes (Figure S8A). In addition, co-culture with brain pericyte-like cells also reduced accumulation and transcytosis of 10 kDa dextran in primary rat BMECs (Figure S8B-C). In summary, these data indicate that hPSC-derived brain pericyte-like cells can induce BBB phenotypes including elevation of BMEC barrier tightness and reduction in transcytosis.

### iPSC-derived brain pericyte-like cells can be integrated into an isogenic NVU model

Previously, we demonstrated that sequential co-culture of iPSC-derived BMECs with primary pericytes and primary neural progenitor-derived astrocytes and neurons enhanced BMEC barrier tightness (*30*). Subsequently, iPSC-derived astrocytes and neurons were shown to induce barrier formation in iPSC-derived BMECs (*34*). Here, iPSC-derived brain pericyte-like cells were combined with iPSC-derived BMECs, astrocytes, and neurons to model the NVU. IMR90C4-derived BMECs were sequentially co-cultured with IMR90C4-derived brain pericyte-like cells and IMR90C4-derived astrocyte/neuron cultures (PNA) and compared to IMR90C4-derived BMEC monocultures or IMR90C4-derived BMECs co-cultured with pericytes (P) or astrocytes/neurons (NA) alone (Figure 6A). All three co-culture conditions (P, NA, and PNA) significantly elevated TEER above monoculture (Figure 6B). While neuron/astrocyte co-culture slightly elevated TEER above pericyte co-culture (720 ± 84 Ω·cm^2^ NA co-culture vs. 503 ± 63 Ω·cm^2^ P co-culture), the combination of pericyte and neuron/astrocyte co-culture treatments further elevated BMEC TEER (1156 ± 94 Ω·cm^2^ vs. NA and P co-culture) (Figure 6B). All three co-culture conditions yielded a five-fold reduction in sodium fluorescein permeability compared to monoculture conditions but no appreciable differences were observed between separate co-culture treatment conditions (Figure 6C), as has been reported previously for BMEC monolayers with TEER values exceeding ~500–600 Ω·cm^2^ (*30, 34, 56*). These data demonstrate that iPSC-derived brain pericyte-like cells can be readily combined with iPSC-derived BMECs, astrocytes and neurons to form an isogenic model of the human NVU.

**Figure 6:**
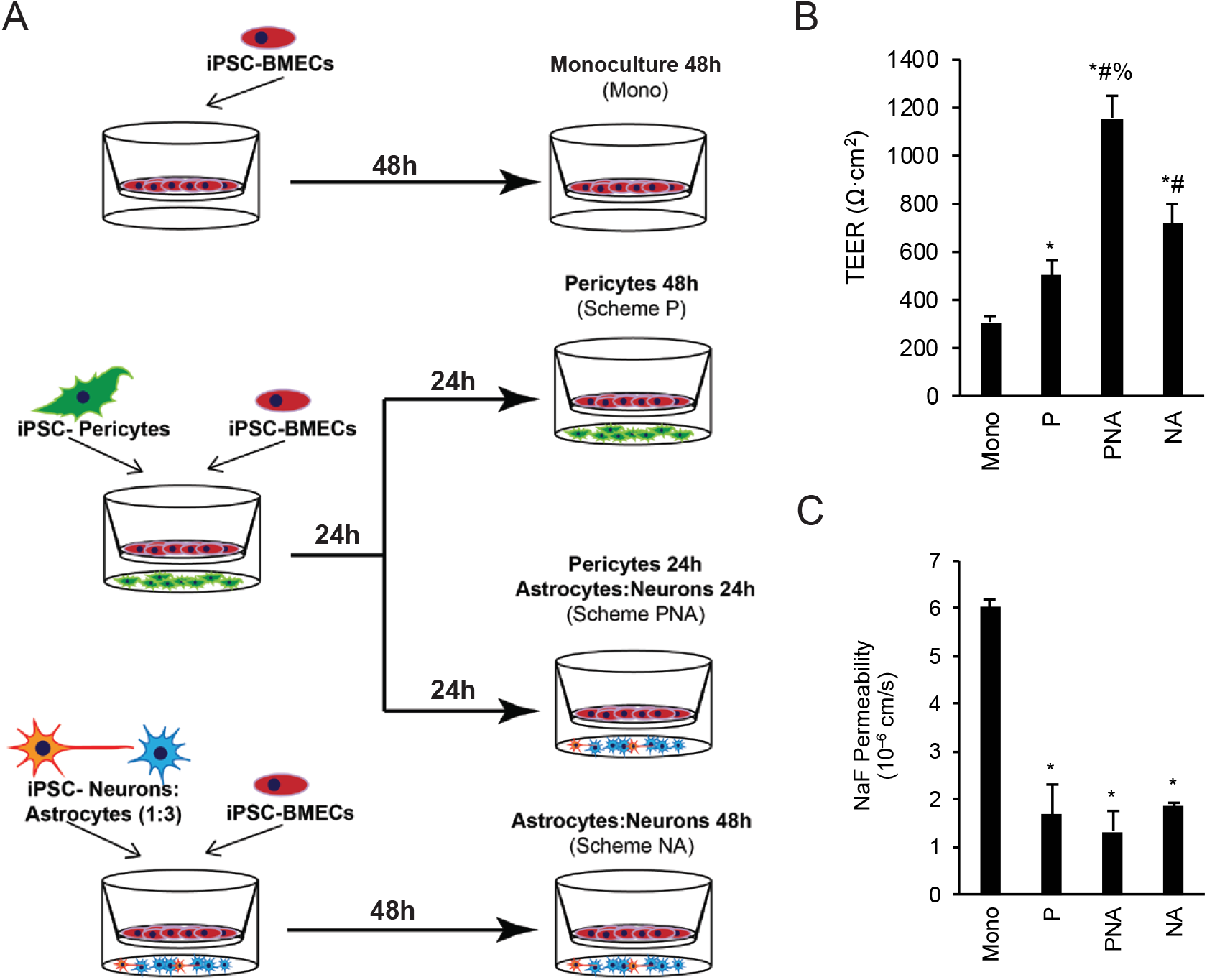
IMR90C4-derived pericyte-like cells integrate into a complete isogenic NVU model. A) Schematic of IMR90C4-derived BMEC co-culture set up with IMR90C4-derived NVU cell types. B) Maximum TEER achieved in IMR90C4-derived BMECs following monoculture or co-culture. Plotted are the means ± SD from 3 Transwells. Results are representative of 3 independent differentiations. * *P* < 0.05 vs. monoculture; # *P* < 0.05 vs. pericyte-like cell co-culture; % *P* < 0.05 vs. astrocyte/neuron co-culture; ANOVA followed by Tukey’s HSD test. C) Sodium fluorescein permeability in IMR90C4-derived BMECs following 48-hours of monoculture or co-culture. Plotted are the means ± SD from 3 Transwells. Results are representative of 3 independent differentiations. * *P* < 0.05 vs. monoculture; ANOVA followed by Tukey’s HSD test.

## DISCUSSION

Brain pericytes play essential roles in BBB formation and maintenance by regulating BMEC transcytosis, barrier fidelity, vascular structure and stability (*5, 13, 14, 19–21*). Here we report that mural cells can be differentiated from hPSC-derived NCSCs, and that these cells develop brain pericyte-like attributes. The brain pericyte-like cells can self assemble with endothelial cells *in vitro* and impact their vascular network structure. Moreover, the brain pericyte-like cells induce BBB properties, including barrier tightening and reduction of transcytosis in BMECs. Finally, these cells can be incorporated into an isogenic iPSC-derived NVU model, with potential applications in patient-specific NVU modeling.

During embryonic development, NCSCs are first specified at the interface between the neural plate and non-neural ectoderm, and subsequently reside in the dorsal neural tube before migrating throughout the embryo and differentiating to diverse cell types (*57*). Previous NCSC differentiation protocols have relied on differentiating hPSCs to neuroectoderm and subsequently isolating NCSC subpopulations (*40, 46*), or have used a directed WNT activation and activin/nodal inhibition approach to obtain NCSCs (*45, 58, 59*). We chose to utilize the latter approach given its simplicity and potential for highly enriched NCSC populations. BMP signaling activation was previously shown to inhibit NCSC formation (*45*); however the need to inhibit BMP signaling during NCSC differentiation has been variable (*41, 45*). Here, when the differentiation strategy was adapted to minimal E6 medium, inhibition of BMP signaling was necessary to efficiently direct hPSCs to p75-NGFR^+^/HNK1^+^ NCSCs. The NCSCs differentiated in E6-CSFD medium were a highly enriched population of multipotent cells having the capacity to form mesenchymal derivatives and peripheral neurons using multiple hPSC lines.

A common approach to differentiate mural cells from NCSCs is to supplement basal medium with PDGF-BB and TGFβ1 (*40, 41, 46*). Resultant cells have been shown to express calponin, SM22α and α-SMA (*40, 41, 46*), but two key mural cell markers, PDGFRβ and NG2 were not previously examined. While differentiation of NCSCs in E6 medium yielded PDGFRβ+ cells, neither E6 medium nor E6 medium supplemented with PDGF-BB and TGFβ1 generated cells expressing NG2. However, both calponin and SM22α were expressed even in the absence of growth factor supplementation. Instead, when E6 was supplemented with 10% FBS, the differentiating cells acquired NG2 and PDGFRβ expression, and thus were classified as forebrain lineage mural cells (*39, 48*). While others have suggested the use of serum to drive mural cell differentiation from NCSC (*45, 46*), these studies generated cells that were smooth muscle actin positive. In contrast, we did not observe substantial α-SMA expression in the differentiated mural cells, even after extended culture. Brain pericytes lining higher order capillaries generally do not express α-SMA *in vivo* (*39, 60, 67*), although very recent evidence suggests that higher order pericytes may actually express low levels of α-SMA that are lost upon sample preparation (*62*). In addition, it is well known that upon fresh isolation, primary brain pericytes express α-SMA in 5-10% of cells, whereas after a few days in culture they become nearly uniformly α-SMA+ (*37, 38*), as also observed here with primary human brain pericytes, which expressed α-SMA. Thus, the lack of α-SMA expression in the differentiated brain pericyte-like cells better reflects the lack of α-SMA in brain pericytes *in vivo*. However, much like primary brain pericytes and previous reports with NCSC derived mural cells, we observed sustained expression of the contractile-related proteins calponin and SM22α. In addition, differentiation of mesenchymoangioblasts towards pericyte lineages yielded cells that expressed differential levels of calponin (*63*). Although SM22α is an early developmental marker of mural cells (*64*), a recent single cell transcriptomics study strongly suggests that murine brain capillary pericytes *in vivo* do not express calponin-encoding *Cnn1* or SM22α-encoding *Tagln* (*39*). Additional transcript evaluation confirmed the brain signature of the pericyte-like cells (*ZIC1, FOXF2*) (*53, 54*). The brain pericyte-like cells also expressed transcripts for *ABCC9* and *KCNJ8*, two additional markers that differentiate brain capillary pericytes from other mural cell types (*39, 48*), and these markers were further elevated over extended culture times. RNA-sequencing also indicated a transcriptome-wide similarity to primary human brain pericytes and expression of many genes identified as pericyte-enriched by single cell RNA-sequencing in mouse (*39*). Taken together, the hPSC-derived brain-pericyte like cells had marker profiles that suggested the generation of cells similar to brain pericytes.

While the marker expression suggested that the differentiation process generated brain pericyte-like cells, it is most important that the cells recapitulate key functional attributes of brain pericytes. When cultured with HUVECs, brain pericyte-like cells aligned along vascular tubes and extended cell processes. Primary pericytes can reduce endothelial cell tube formation *in vitro* (*26*), and this phenotype was also observed with both primary human brain pericytes and hPSC-derived brain pericyte-like cell co-culture as indicated by reduced number of segments per field. Also, the length of tube segments was increased in the presence of primary or hPSC-derived brain pericytes as has been reported previously with hPSC-derived pericytes of mesenchymal origins (*63*). In addition to this more general pericyte phenotype, it was expected that a brain pericyte-like cell would impact the barrier and transcytosis properties of brain endothelial cells (*13, 14*). Indeed, BBB properties of both hPSC-derived BMECs and primary rat BMECs were substantially induced by co-culture with hPSC-derived brain pericyte-like cells, and these effects mimicked those induced by primary human brain pericytes. TEER was increased substantially as expected (*13, 30*). Correlating with this increased barrier function, pericyte co-culture decreased the number of frayed tight junctions as seen previously for a variety of barrier inductive stimuli^80,82,96^, but did not alter the expression levels of tight junction proteins occludin or claudin-5. These results mirror those *in vivo* where tight junction structure was altered by pericytes although the expression levels of tight junction proteins were not affected (*13*). We also demonstrated that nonspecific cellular accumulation and transcytosis were downregulated in BMECs after co-culture with brain-pericyte like cells, and the effects were indistinguishable from those elicited by primary human brain pericytes. These phenotypes combined with the developmental origins and marker expression profile, along with the similarities to primary human brain pericytes, suggest that we have generated a novel hPSC-derived cell that can model human brain pericytes.

While many studies have utilized primary pericytes to enhance BMEC barrier properties, primary pericytes offer limited scalability, especially for human *in vitro* BBB models (*28, 30, 65*). In addition, limited primary cell availability essentially eliminates the possibility of using patient matched brain pericytes and BMECs that could be used for disease modeling applications. Here, we demonstrate the capability to differentiate brain like pericytes in a scalable fashion (~1000 brain pericyte-like cells per input stem cell). Moreover, the differentiation is reproducible amongst differentiations and across hPSC lines. Thus, the ability to derive brain pericyte-like cells from patient-derived iPSCs provides a unique tool for the study of patient-specific pericyte contributions to CNS disorders that have been suggested to have pericyte involvement such as stroke, epilepsy, demyelinating disease, and Alzheimer’s disease (*7, 10, 66–69*). In addition, lineage-specific differences have been noted in hPSC-derived pericytes, motivating the use of pericytes from appropriate developmental origins for disease modeling applications (*40*). The brain pericyte-like cells can also be used in multicellular NVU models to capture the cellular crosstalk that is likely responsible for many disease processes at the BBB. To this end, we have demonstrated that it is possible to generate BMECs, neurons, astrocytes, and brain pericyte-like cells from a single iPSC cell line and combine them to form an isogenic NVU model having optimal TEER. These findings followed similar trends to our earlier reports where iPSC-derived BMEC properties were enhanced by co-culture with brain pericytes and neural progenitor cell-derived astrocytes and neurons from primary sources. It is likely that these multicellular NVU models will be used to uncover new mechanisms of BBB regulation in health and disease and assist in the therapeutic development process for CNS disorders.

## MATERIALS AND METHODS

### hPSC maintenance

IMR90C4 and CS03n2 iPSCs and H9 hESCs were maintained on Matrigel coated plates in E8 medium containing DMEM/F12, L-ascorbic acid-2-phosphate magnesium (64 mg/L), sodium selenium (14 μg/L), FGF2 (100 μg/L), insulin (19.4 mg/L), NaHCO_3_ (543 mg/L), transferrin (10.7 mg/L), and TGFβ1 (2 μg/L) and prepared according to Chen *et al.* (*70*). When cells reached ~70% confluence, cells were passaged using Versene to new Matrigel coated plates. For hPSCs used in BMEC differentiations, cells were maintained in mTeSR1 on Matrigel plates and passaged as previously described (*71*).

### NCSC differentiation

One day prior to initiating NCSC differentiation, hPSCs maintained in E8 medium were singularized using Accutase and seeded at 9.1 × 10^4^ cells/cm^2^ onto Matrigel coated plates with E8 + 10 μM Y27632. NCSC differentiation was initiated the next day by switching medium to E6, containing DMEM/F12, L-ascorbic acid-2-phosphate magnesium (64 mg/L), sodium selenium (14 μg/L), insulin (19.4 mg/L), NaHCO_3_ (543 mg/L), and transferrin (10.7 mg/L). E6 was supplemented with 22.5 mg/L heparin sodium salt from porcine mucosa, 1 μM CHIR99021, 10 μM SB431542 (Tocris), 10 μg/L FGF2, and 1 μM dorsomorphin, hereafter labeled E6-CSFD. Cells were expanded by replacing E6-CSFD daily and passaging cells every time cells reached 100% confluence to fresh Matrigel coated plates. During passaging, cells were singularized using Accutase and replated at a splitting density of one 6-well to 6 new 6-wells in E6-CSFD medium. Cells were generally passaged without 10 μM Y27632. However, to increase IMR90C4 cell line survival during first passaging following NCSC differentiation initiation, IMR90C4 cells were replated in E6-CSFD + 10 μM Y27632. Subsequent IMR90C4 NCSC expansion passages were replated without Y27632. Cells were typically passaged 2-3 days following NCSC differentiation initiation and subsequently passaged every 3-6 days depending on cell growth kinetics.

### Magnetic activated cell sorting of NCSCs

At D15 of E6-CSFD treatment, cells were dissociated using Accutase and labeled with 20 μL/10^7^ cells NCSC microbeads (Miltenyi), 20 μL/10^7^ cells FcR blocking reagent, and 60 μL/10^7^ MACS buffer (0.5% BSA + 2 mM EDTA in sterile PBS without Ca^2+^/Mg^2+^) at 4°C for 15 minutes. Cells were washed in MACS buffer and resuspended in 500 μL MACS buffer/2× 10^7^ cells. Cells were sorted through two LS columns (Miltenyi) according to manufacturer instructions and resuspended in E6-CSFD + 10 μM Y27632 to appropriate density for specific NCSC lineage differentiations as described below.

### NCSC lineage differentiations

For differentiation of peripheral neurons, after MACS sorting, hPSC-derived NCSCs were replated on Matrigel-coated plates and expanded for 14 days in E6-CSFD. These cells were replated on Matrigel-coated 12-well plates at 5×10^4^ cells/cm^2^ in E6-CSFD. The following day, the medium was switched to peripheral neuron medium composed of DMEM/F12, 1× N2 supplement, 10 ng/ml BDNF, 10 ng/ml GDNF, 10 ng/ml NT-3, 10 ng/ml NGF-β, 200 μM ascorbic acid (AA), and 0.5 mM cAMP, and replaced every 2 days for 2 weeks.

For differentiation of mesenchymal derivatives, after MACS sorting, hPSC-derived NCSCs were replated on noncoated polystyrene plates and expanded for 11 days in E6-CSFD. For adipogenesis, expanded hPSC-derived NCSCs were seeded at a density of 10,000 cells/cm^2^ and treated with adipogenic medium composed of high-glucose DMEM, 10% FBS, 1% antibiotics, 1 μg/ml insulin, 0.5 mM 3-isobutyl-1-methylxanthine (IBMX), and 1 μM dexamethasone (Sigma-Aldrich). For osteogenesis, the seeding density was 5,000 cells/cm^2^ and the cells were treated with osteogenic medium consisting of low-glucose DMEM, 10% FBS, 1% antibiotics, 50 μg/ml AA, 10 mM β-glycerophosphate, and 0.1 μM dexamethasone. For chondrogenesis, 250,000 NCSC were collected to form a high cell density pellet by centrifuged at 600 g for 5 minutes and treated with chondrogenic medium containing high-glucose DMEM, 1% antibiotics and ITS Premix (BD Bioscience), 40 μg/ml L-proline, 50 μg/ml AA, 0.9 mM sodium pyruvate (Sigma-Aldrich), 0.1 μM dexamethasone, and 10 ng/ml of freshly added transforming growth factor β 1 (TGFβ1) (Peprotech). Medium was changed every 3 days for all three differentiation procedures.

To analyze adipogenic differentiation, cells were fixed in 4% of formaldehyde and stained with Oil Red O (Sigma-Aldrich) for lipid droplet formation. To analyze osteogenic differentiation, cells were fixed in 60% isopropanol and stained with Alizarin Red (Rowley Biochemical, Danvers, MA, USA) to evaluate mineral deposition. Chondrogenic potential was assessed by Alcian blue staining. Cell pellets were first fixed in 4% formaldehyde for 2 hours. Next, the cell pellet was dehydrated by a series of increasing concentration of ethanol, infiltrated with xylene, and then embedded with paraffin. Embedded cell pellets were cut into 8 μm sections using a microtome and stained with Alcian blue (Polysciences, Warrington, PA, USA) to determine the glycosaminoglycan (GAG) content.

### Pericyte differentiation factor identification

Following MACS sorting, NCSC were replated onto 48-well plates in E6-CSFD medium + 10 μM Y27632. Cells were switched to mural cell differentiation medium the next day, expanded for six days, and stained for NG2/PDGFRβ expression. Cells were expanded on uncoated plates in E6 medium, E6 medium supplemented with 2 ng/mL TGFβ1 + 20 ng/mL PDGF-BB, or E6 medium supplemented with 10% fetal bovine serum (FBS). Cells were also expanded in E6 supplemented with 10% FBS on gelatin-coated plates prepared by coating plates for at least 1h at 37°C with a 0.1% gelatin A solution dissolved in water.

### Immunocytochemistry

Cells were fixed fifteen minutes at room temperature with either 4% paraformaldehyde (PFA) or 100% ice-cold methanol depending on antibody staining conditions. Cells were rinsed three times in PBS without Ca^2^+/Mg^2^+ and stored at 4°C in PBS until ready to stain. After aspirating PBS, cells were blocked one hour in blocking buffer at room temperature and incubated overnight at 4°C on a rocking platform with primary antibodies diluted in primary antibody staining buffer. Antibodies and staining conditions are listed in Table S1. The following day, cells were washed three times with PBS and incubated with secondary antibodies diluted 1:200 in primary antibody staining buffer. Cells were probed one hour in the dark at room temperature on a rocking platform. Afterwards, secondary antibody staining buffer was aspirated, and cells were incubated five minutes with 4 μM Hoechst 33342 diluted in PBS. Cells were washed three times with PBS and stored at 4°C in PBS in the dark until ready to image. Images were taken on Olympus epifluorescence and Nikon A1R-Si+ confocal microscopes.

### Flow cytometry

Cells were incubated 30 minutes on ice with primary antibody diluted in 100 μL/sample primary antibody staining buffer as indicated in Table S1. Cells were washed one time with cold PBS (p75-NGFR/HNK1 flow cytometry) or MACS buffer (NG2 and PDGFRβ flow cytometry). Cells were subsequently incubated in 100 μL primary antibody staining buffer with 1:500 Alexa-tagged isotype-specific goat secondary antibodies. Cells were washed as previously described and resuspended in 4% PFA for 15 minutes at room temperature. Cells were subsequently stored in wash buffer for up to 24 hours at 4°C prior to running samples on cytometer.

### Temporal RNA analysis

Cells were harvested using Accutase, quenched in DMEM/F12, and spun down 5 minutes at 200g. After removing the supernatant, cell pellets were snap frozen at −80°C until ready for mRNA extraction. The RNeasy Mini Kit (Qiagen) was used to extract mRNA, including a cell lysate homogenization step on QIAshredder Columns (Qiagen), according to manufacturer instructions. DNA was removed on column using the RNase-free DNase Set (Qiagen). Extracted RNA was stored in nuclease-free water at −20°C until ready to reverse transcribe to cDNA. RNA was reverse transcribed at a concentration of 250 ng/mL into cDNA using Omniscript reverse transcriptase kit (Qiagen) and Oligo(dT)_20_ Primers (Life Technologies). Temporal gene expression analysis was conducted using 25 μL PCR reactions containing GoTaq Green Master Mix (Promega), 10 ng/reaction cDNA template, and 100 nM forward/reverse primers. PCR was run according to manufacturer protocols, and all reactions included a no template and mRNA control to verify no genomic DNA contamination or amplification. PCR primer sequences, annealing temperatures, and cycle times are listed in Table S2. PCR products were resolved on a 2% agarose gel, stained using ethidium bromide, and imaged on a ChemiDoc XRS+ System (Bio-Rad).

### RNA-sequencing

RNA was extracted from H9 hESCs, H9-derived NCSCs at D15, H9-derived NCSCs maintained for 40 additional days in E6-CSFD (D55), H9-derived pericyte-like cells at D19,D22, and D25 (three independent differentiations at the D25 time point), H9-derived pericyte-like cells maintained for 20 additional days in E6 + 10% FBS (D45), CS03n2-derived pericyte-like cells at D25, IMR90C4-derived pericyte-like cells at D25, and primary brain pericytes using the RNeasy Mini Kit (Qiagen) as described above. TruSeq stranded mRNA libraries were prepared, cDNA synthesized, pooled, and distributed over two sequencing lanes, and samples sequenced on an Illumina HiSeq 2500 by the University of Wisconsin-Madison Biotechnology Center. Reads were mapped to the human genome (hg38) with HISAT2 (v2.1.0) and transcript abundances (fragments per kilobase of transcript per million mapped reads, FPKM) quantified with Cufflinks (v2.1.1). FPKM values from the two sequencing lanes for each sample were averaged. Hierarchical clustering was performed with Morpheus (https://software.broadinstitute.org/morpheus) using the one minus Pearson correlation with average linkage. Gene ontology (GO) analysis was performed using the PANTHER (*72*) online tool (http://pantherdb.org).

### Matrigel tube formation assay and quantification

8-well glass chamber slides were coated with 200 μL/well concentrated growth factor reduced Matrigel and incubated at least one hour at 37°C to set the Matrigel. Human umbilical vein endothelial cells (HUVECs) were plated at 2.2×10^4^ cells/8-well chamber slide in 500 μL EGM-2 medium (Lonza) alone, with 6.6×10^4^ HEK293 fibroblasts, primary brain pericytes, or hPSC-derived mural cells at D22 of the differentiation. Cells were incubated 24 hours at 37°C to allow tube formation and bright field images taken on live cells at 24 hours following plating. Tubes were subsequently fixed and stained according the immunocytochemistry methods listed above. Matrigel-associated tubes were mounted onto glass slips and imaged using Olympus epifluorescence and Nikon A1R-Si+ confocal microscopes. Tube length and number of tubes per field were quantified by hand using the ImageJ ROI Manager Tool and averaged over at least 3 independent fields per condition per differentiation.

### BMEC differentiation

IMR90C4 iPSCs were maintained in mTesR1 medium on Matrigel-coated plates and passaged as previously described. Three days prior to initiating a differentiation, cells were seeded at 9×10^4^ – 10^5^ cells/cm^2^ onto Matrigel coated plates in mTeSR1 + 10 μM Y27632. Medium was replaced daily until cells reached > 2.6×10^5^ cells/cm^2^. Cell medium was replaced with Unconditioned Medium (UM), containing 392.5 mL DMEM/F12, 100 mL KOSR (Gibco), 5 mL 100× MEM non-essential amino acids (Gibco), 2.5 mL 100× Glutamax (Gibco), and 3.5 μL β-mercaptoethanol. Cells were replaced daily with UM for six days, and subsequently switched to EC medium, containing hESFM + 1% platelet-poor plasma derived serum (PDS), and 20 ng/mL FGF2. Cells were incubated two days with EC medium without replacing medium. Cells were sub-cultured at D8 onto 4:1:5 collagen/fibronectin/water-coated Transwells or 5× diluted 4:1:5 collagen/fibronectin/water-coated plates as detailed by Stebbins *et al.* (*71*). Cell culture medium was replaced with EC without FGF2 24 hours after subculturing hPSC-derived BMECs onto filters.

### BMEC/pericyte co-culture

Primary brain pericytes, hPSC-derived pericyte-like cells, and 3T3s were separately seeded onto poly-L-lysine coated 12-well plates (primary brain pericytes) or uncoated plates (hPSC-derived early mural cells and 3T3s) when early mural cells reached first reached 80-100% confluence, typically 3-4 days after initiating serum treatment on hPSC-derived NCSC (D19-D20). Cells were plated on the same day at equivalent seeding densities of 5×10^4^ cells/12-well in either DMEM + 10% FBS (primary brain pericytes and 3T3s) or E6 + 10% FBS (hPSC-derived pericyte-like cells). Cells were dissociated with either 0.25% Trypsin/EDTA (primary brain pericytes and 3T3s) or Accutase (hPSC-derived pericyte-like cells). hPSC-derived brain pericyte-like cell medium was replaced daily with E6 + 10% FBS until D22 of the differentiation. Pericytes and 3T3s were fed with DMEM + 10% FBS every two days until D22. At D22, cells were replaced with 1.5 mL EC medium above 12-well polystyrene transwell filters with a 0.4 μm pore size.

IMR90C4-derived BMECs at D8 of the BMEC differentiation were sub-cultured onto Transwell filters in co-culture with BMECs at 1.1×10^6^ cells/12-well filter as previously described. Cells were incubated two days in co-culture, with cell culture medium replaced at 24 hours after initiating co-culture with EC medium without FGF2. Transendothelial electrical resistance (TEER) was measured every 24 hours after initiating co-culture. 48 hours following co-culture, BMEC Transwell filters were transferred to a fresh 12-well plate for sodium fluorescein assays. Cell medium was replaced with 1.5 mL of EC medium without FGF2 in the basolateral chamber and 0.5 mL of the same medium with 10 μM sodium fluorescein in the apical chamber. Cells were incubated one hour on a rotating platform and basolateral chamber medium collected every 15 minutes during the hour incubation period. After 1 hour, cell culture medium for the apical chamber was collected to calculate sodium fluorescein permeability across BMEC monolayers following 48 hours of co-culture treatment. Fluorescence intensity was measured using a Tecan plate reader set to a 485 nm excitation and 530 nm emission settings. Permeability calculations were determined according to Stebbins *et al.* (*71*).

### Transcytosis and accumulation assays

Following 48 hours of co-culture, BMEC-seeded transwells were transferred to an empty plate. We utilized a 10 kDa dextran tagged with Alexa-488 to quantify the level of intracellular accumulation and transcytosis. 10 μM dextran was suspended in 0.5 mL of EC medium without FGF2 onto the apical side of the transwell. To determine the level of transcytosis, following two hours of incubation in a 37°C incubator (20% O_2_, 5% CO_2_) on a rotating platform, we collected 150 μL from the 1.5 mL of EC medium on the basolateral side of the transwell. Fluorescence intensity was measured using a Tecan plate reader set to a 495 nm excitation and 519 nm emission settings. To determine the level of accumulation (endocytosis) in the BMECs, we rinsed the transwells with cold PBS (2×) and lysed the cells with radioimmunoprecipitation assay (RIPA) buffer. Lysates were collected and analyzed on the plate reader. Fluorescence values were normalized to protein content/Transwell measured using the bicinchoninic acid (BCA) assay.

### Tight junction image analysis

BMECs were plated on 24-well plates in EC medium or EC medium conditioned by primary brain pericytes or hPSC-derived pericyte-like cells. After 48 h, BMECs were fixed and stained for occludin as described above. Images were acquired from 5 wells per experimental condition. To quantify tight junction continuity, images were blinded and the area fraction index determined using FIJI as previously described (*73*). Additionally, images were blinded and the number of frayed junctions (Figure 4D) manually counted.

### Isogenic model of the neurovascular unit

BMECs were differentiated from iPSCs as previously described. Singularized BMECs were seeded onto collagen IV/fibronectin coated transwells at day 8 of the differentiation. hPSC-derived pericytes were seeded onto the bottom of the co-culture plate (~50,000 cells/cm^2^) in EC medium. We additionally investigated the cumulative effects of pericytes, neurons, and astrocytes. Neurons and astrocytes were differentiated from iPSCs as previously published (*34*). Initially, BMECs were placed in co-culture with pericytes for 24 hours in EC medium and then BMEC-Transwells were transferred to a co-culture plate with a mixture of neurons and astrocytes (1:3 ratio) for the duration of the experiment in EC medium without FGF2. We benchmarked our stem cell-derived BBB model (BMECs, pericyte-like cells (24h), and neurons and astrocytes (24h)) to a BBB co-culture model absent of pericyte-like cells (neurons and astrocytes only). TEER and sodium fluorescein permeability assays were conducted on BMEC-Transwells.

### Statistics

All experiments were conducted using at least three technical replicates (e.g. three 6- wells or Transwells) from the same differentiation. All experiments were replicated (independent differentiations) at least three times except where otherwise indicated. Data are presented as mean ± SD of technical replicates from a representative differentiation or as mean ± SEM of pooled data from several independent differentiations. Statistical significance was evaluated using one-way analysis of variance (ANOVA) followed by post-hoc tests controlling for multiple comparisons: Dunnett’s test for comparison of experimental groups to control, and Tukey’s test for comparison between all experimental groups. *P* < 0.05 was considered statistically significant.

### Western blotting

BMECs were cultured on Transwells alone or co-cultured with hPSC-derived pericyte-like cells, primary brain pericytes, or 3T3s as previously described. After 48 h of co-culture, BMECs were washed once with PBS and lysed with RIPA buffer + Halt protease inhibitor cocktail. The BCA assay was used to determine protein concentration. Proteins were resolved on 4-12% Tris-glycine gels and transferred to nitrocellulose membranes, which were blocked in Tris-buffered saline + 0.1% Tween-20 (TBST) + 5% nonfat dry milk for 1 h, and incubated with primary antibodies (Table S1) overnight at 4°C. Membranes were washed with TBST (5×) and incubated with donkey anti-mouse or donkey anti-rabbit IRDye 800CW secondary antibodies (LI-COR) for 1 h. Membranes were washed with TBST (5×) and imaged using a LI-COR Odyssey.

### Visualization of dextran accumulation

iPSC-derived BMECs were seeded onto glass bottom plates at a density of 10^5^ cells/cm^2^ and cultured for 48 h in EC medium or EC medium conditioned by 3T3s, primary pericytes, or IMR90C4-derived pericyte-like cells. Medium was subsequently replaced with EC medium + 10 μM Alexa 488-tagged 10 kDa dextran. Following 2 h of dextran incubation, cells were fixed in 4% PFA for fifteen minutes, followed by three washes in PBS. Cells were blocked in 10% goat serum in PBS for 30 minutes at room temperature. Cells were incubated with Anti-Alexa 488 antibody (1:100 in PBS; Life Technologies 11094) overnight at 4°C on a rocking platform. Following three washes in PBS, cells were incubated with Alexa 647 secondary antibody (1:200 in PBS) for one hour at room temperature on a rocking platform. Nuclei were labeled with Hoechst and cells were rinsed three times in PBS. Cells were visualized on a Nikon A1R-Si+ confocal microscope. The lack of permeabilization allows internalized dextran to visualized only with Alexa 488, while extracellular (surface) dextran is also labeled with Alexa 647.

### Primary rat BMEC/pericyte co-culture

All animal studies were conducted using protocols approved by the University of Wisconsin-Madison Animal Care and Use Committee following NIH guidelines for care and use of laboratory animals. Adult male Sprague-Dawley rat (Harlan Inc., Indianapolis, IN) brain capillaries were isolated, minced and digested in collagenase type-2 (0.7 mg/mL) and DNase I (39 U/mL). Purified microvessels were isolated following centrifugation in 20% bovine serum albumin and further digested in collagenase/dispase (1 mg/mL) and DNase I. To purify the population a 33% Percoll gradient was utilized. Capillaries were collected and plated onto collagen IV/fibronectin-coated Transwells. Capillaries were cultured in DMEM supplemented with 1 ng/mL FGF2, 1 μg/mL heparin, 20% PDS, 2 mM L-glutamine, and 1% antibiotic-antimitotic solution. Pure populations were obtained by treating the cells with puromycin (4 μg/mL) for 2 days following seeding. Four days following isolation, rat BMEC-seeded Transwells were transferred onto plates containing IMR90C4-derived pericyte-like cells, primary brain pericytes, or 3T3s (described previously) and co-cultured in EC medium containing 1% PDS.

## ACKNOWLEDGMENTS

This work was supported by NIH grants NS083688 and NS103844 (EVS, SPP, and RD) and DTRA grant HDTRA-15-1-0012 (EVS). We would like to acknowledge the University of Wisconsin-Madison Biotechnology Center Gene Expression Center and DNA Sequencing Facility for providing library preparation and next generation sequencing services. MJS and BDG were supported by National Institutes of Health Biotechnology Training Program grant T32 GM008349. BDG was supported by the National Science Foundation Graduate Research Fellowship Program under grant number 1747503.

## AUTHOR CONTRIBUTIONS

MJS, SPP, and EVS conceived the project. MJS, BDG, SGC, M-SL, W-JL, RD, SPP, and EVS designed and analyzed experiments. MJS, BDG, SGC, M-SL, DR, and MGF performed experiments. MJS, BDG, SPP, and EVS wrote the manuscript.

## DATA AVAILABILITY STATEMENT

The data that support the conclusions in the paper are available within the paper or its associated supplementary content files.

---

**Figure S1:**
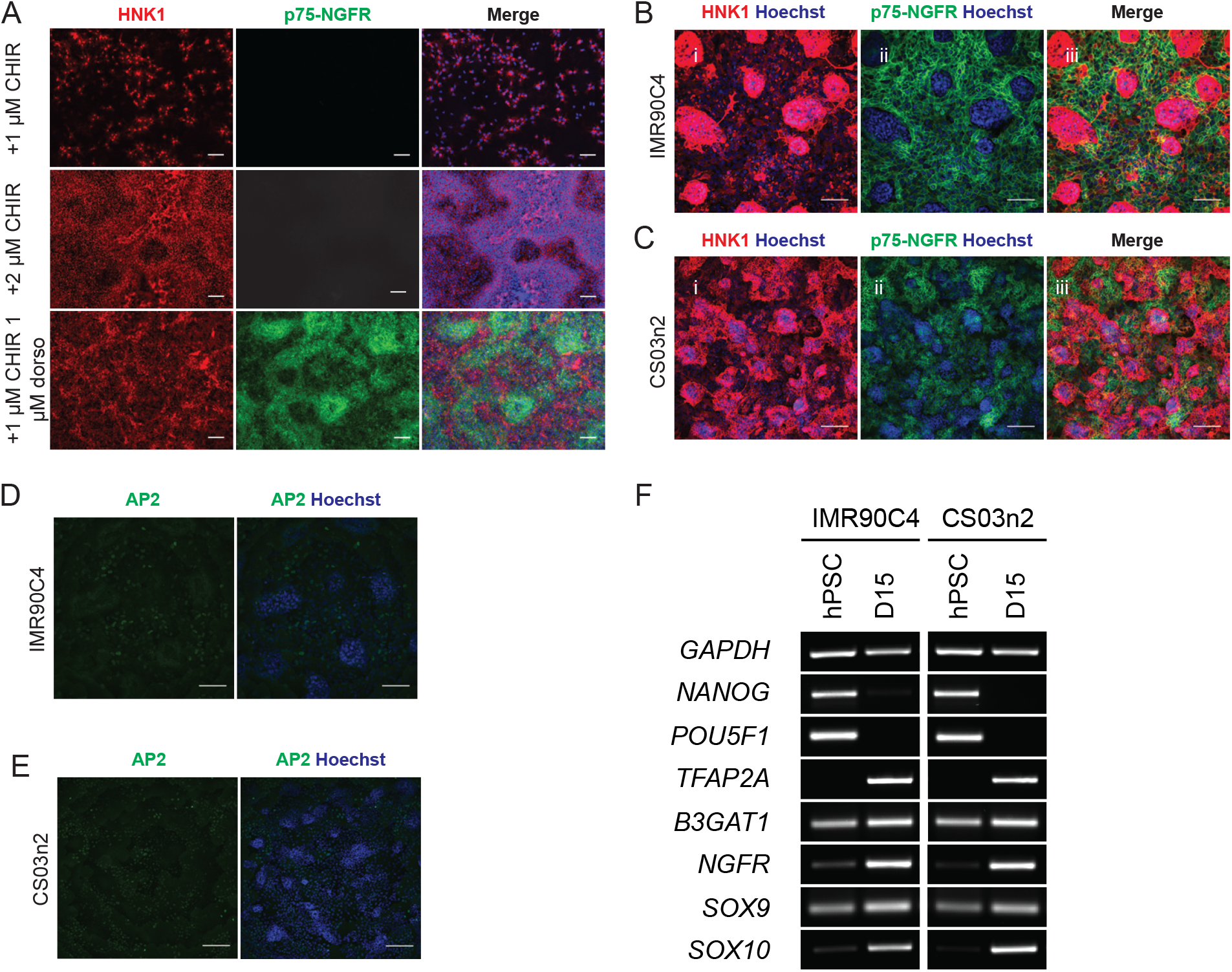
Generation of multipotent NCSC populations from multiple hPSC lines. A) Immunocytochemistry images of small molecule screen (n = 1) on HNK1 and p75-NGFR expression in cells differentiated from H9 hESCs. Cells were cultured fifteen days in E6 + 10 ng/mL FGF2 + 22.5 μg/mL heparin + 10 μM SB431542 + CHIR99012 ± dorsomorphin at the indicated concentrations. Hoechst nuclear counter stain (blue) is also included. Scale bars: 100 μm. B) Immunocytochemistry images of IMR90C4 iPSCs differentiated in E6-CSFD probed for the presence of HNK1 and p75-NGFR at D15. Hoechst nuclear counter stain (blue) is also included. Scale bars: 100 μm. C) Immunocytochemistry images of CS03n2 iPSCs differentiated in E6-CSFD probed for the presence of HNK1 and p75-NGFR at D15. Hoechst nuclear counter stain (blue) is also included. Scale bars: 100 μm. D) AP-2 immunocytochemistry images for IMR90C4-derived NCSCs at D15. Hoechst nuclear counter stain (blue) is also included. Scale bar: 100 μm. E) AP-2 immunocytochemistry images for CS03n2-derived NCSCs at D15. Hoechst DNA nuclear stain (blue) is also included. Scale bar: 100 μm. F) Temporal PCR analysis of pluripotency (NANOG, *POU5F1*) and NCSC (*TFAP2A, B3GAT1, NGFR, SOX9, SOX10*) transcripts in IMR90C4 and CS03n2 iPSCs and NCSC progeny.

**Figure S2:**
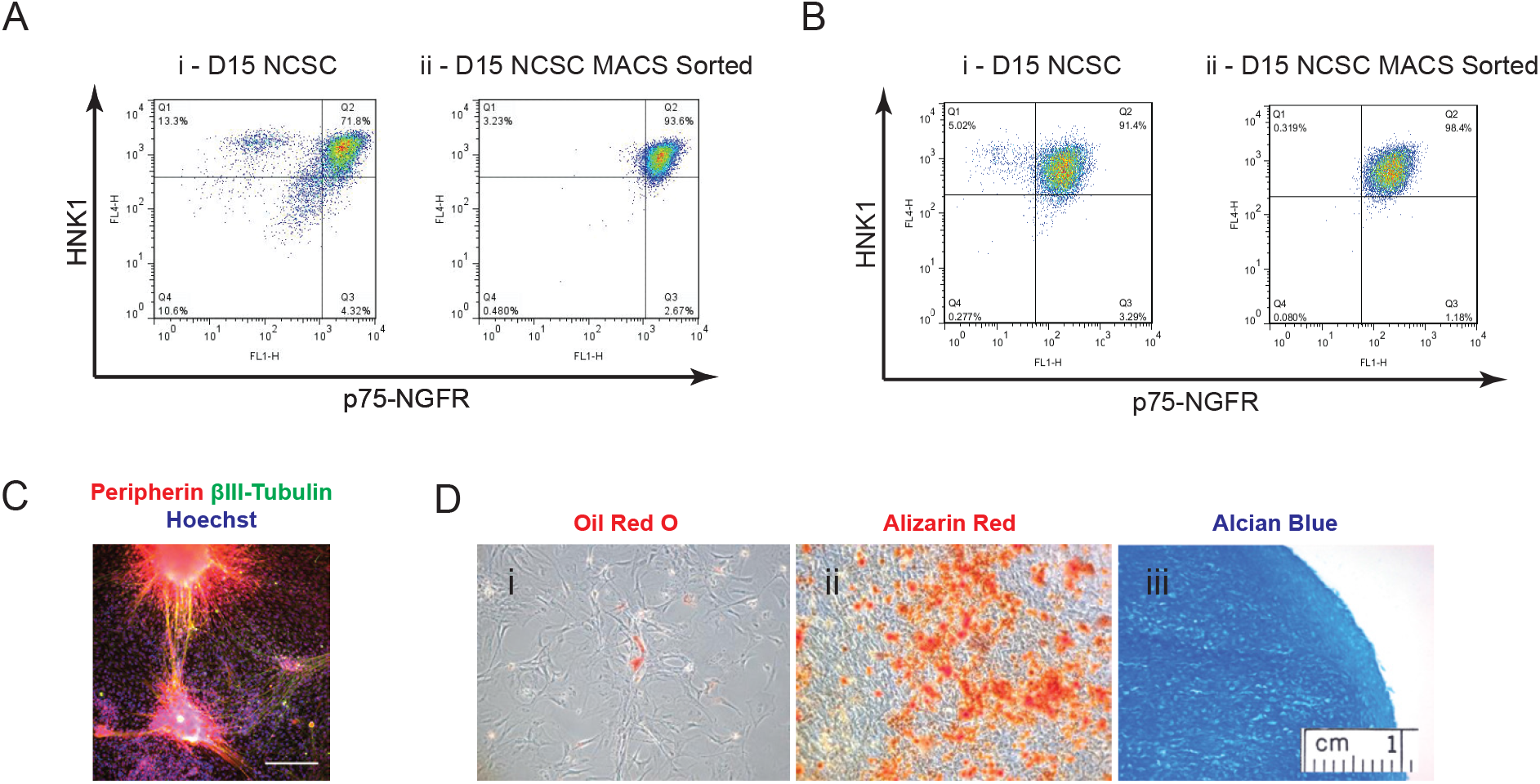
Lineage analysis of iPSC-derived NCSCs. A) Flow cytometry analysis of IMR90C4-derived NCSCs. Panels include NCSC (HNK1^+^/p75-NGFR^+^) purity prior to MACS (panel i), and NCSC purity following MACS (panel ii). Inset percentages are included in each quadrant. Quantitation is shown in Table 1. B) Flow cytometry analysis of CS03n2-derived NCSCs. Panels include NCSC (HNK1+/p75-NGFR+) purity prior to MACS (panel i), and NCSC purity following MACS (panel ii). Inset percentages are included in each quadrant. Quantitation is shown in Table 1. C) Immunocytochemistry analysis of IMR90C4-derived NCSCs subsequently differentiated in peripheral neuron medium. Resultant cells were positive for βIII-tubulin and peripherin expression. Hoechst nuclear counter stain (blue) is also included. Scale bar: 100 μm. D) IMR90C4-derived NCSCs could be differentiated into mesenchymal derivatives, including Oil Red O stained adipocytes (panel i, red), Alizarin red stained osteocytes (panel ii, red), and Alcian blue stained chondrocytes (panel ii, blue).

**Figure S3:**
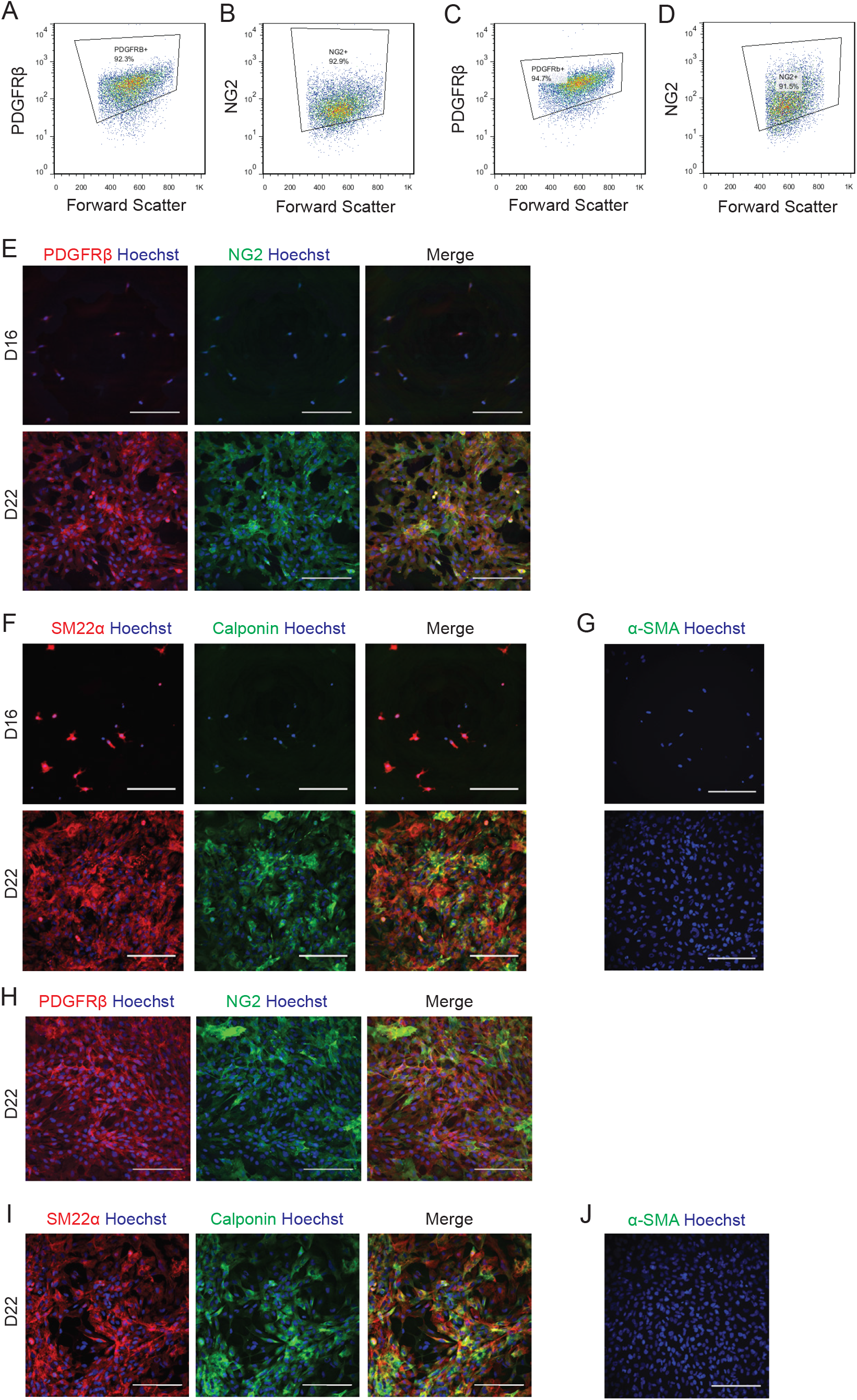
Serum treatment directs iPSC-derived NCSCs towards mural cells. A, B) Representative PDGFRβ and NG2 flow cytometry plots for IMR90C4-derived NCSCs treated for 9 days with E6 + 10% FBS medium. C, D) Representative PDGFRβ and NG2 flow cytometry plots for IMR90C4-derived NCSCs treated for 9 days with E6 + 10% FBS medium. Quantitative results can be found in Table 1. E) PDGFRβ and NG2 immunocytochemistry of IMR90C4-derived NCSCs (D16) and mural cells (D22). Hoechst nuclear counter stain (blue) is also included. Scale bar: 200 μm. F) Calponin and SM22α immunocytochemistry of IMR90C4-derived NCSCs (D16) and mural cells (D22). Hoechst nuclear counter stain (blue) is also included. Scale bar: 200 μm. G) α-SMA immunocytochemistry of IMR90C4-derived NCSCs (D16) and mural cells (D22). Hoechst nuclear counter stain (blue) is also included. Scale bars: 200 μm. H) PDGFRβ and NG2 immunocytochemistry of CS03n2-derived mural cells (D22). Hoechst nuclear counter stain (blue) is also included. Scale bar: 200 μm. I) Calponin and SM22α immunocytochemistry of CS03n2-derived mural cells (D22). Hoechst nuclear counter stain (blue) is also included. Scale bar: 200 μm. J) α-SMA immunocytochemistry of CS03n2-derived mural cells (D22). Hoechst DNA nuclear stain (blue) is also included. Scale bars: 200 μm.

**Figure S4:**
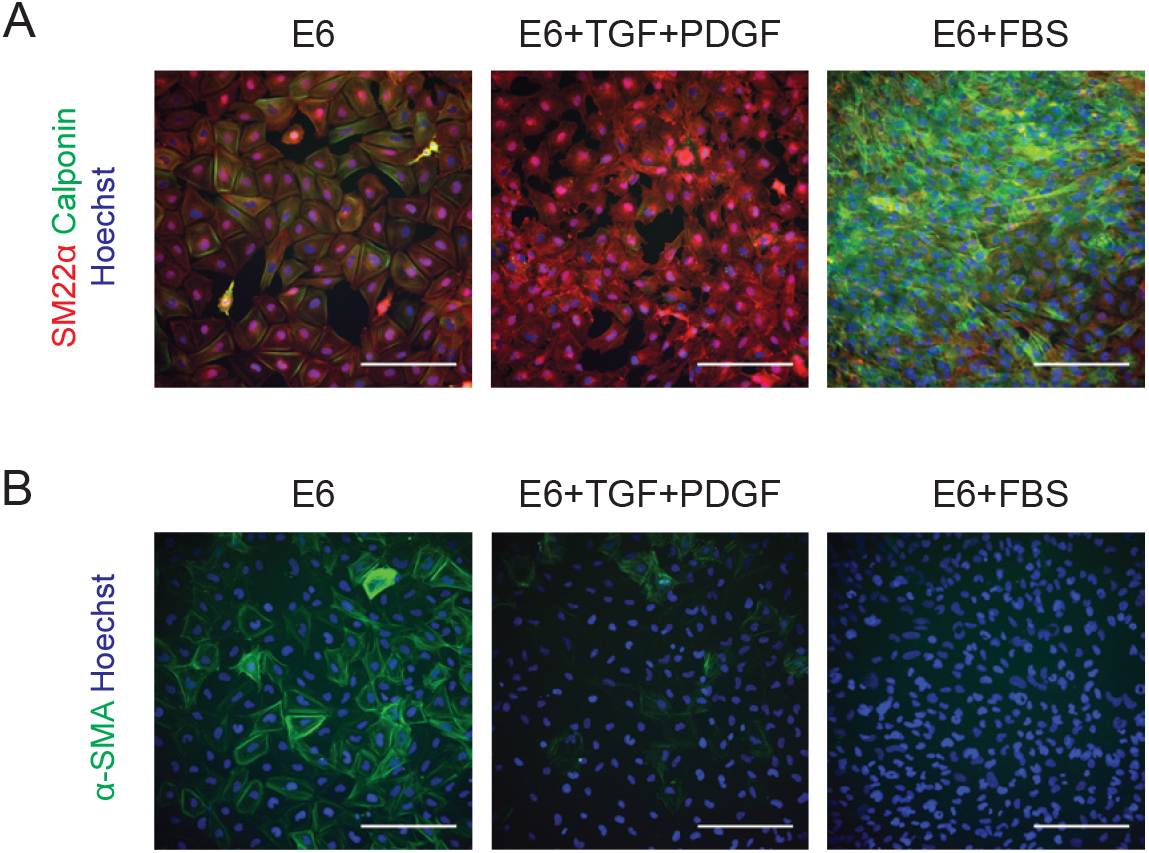
Analysis of cells obtained by culturing NCSCs in E6, E6 + TGFβ1 + PDGF-BB, or E6 + 10% FBS for 6 days. A) Calponin and SM22α immunocytochemistry. Hoechst nuclear counter stain (blue) is also included. Scale bars: 200 μm. B) α-SMA immunocytochemistry. Hoechst nuclear counter stain (blue) is also included. Scale bars: 200 μm.

**Figure S5:**
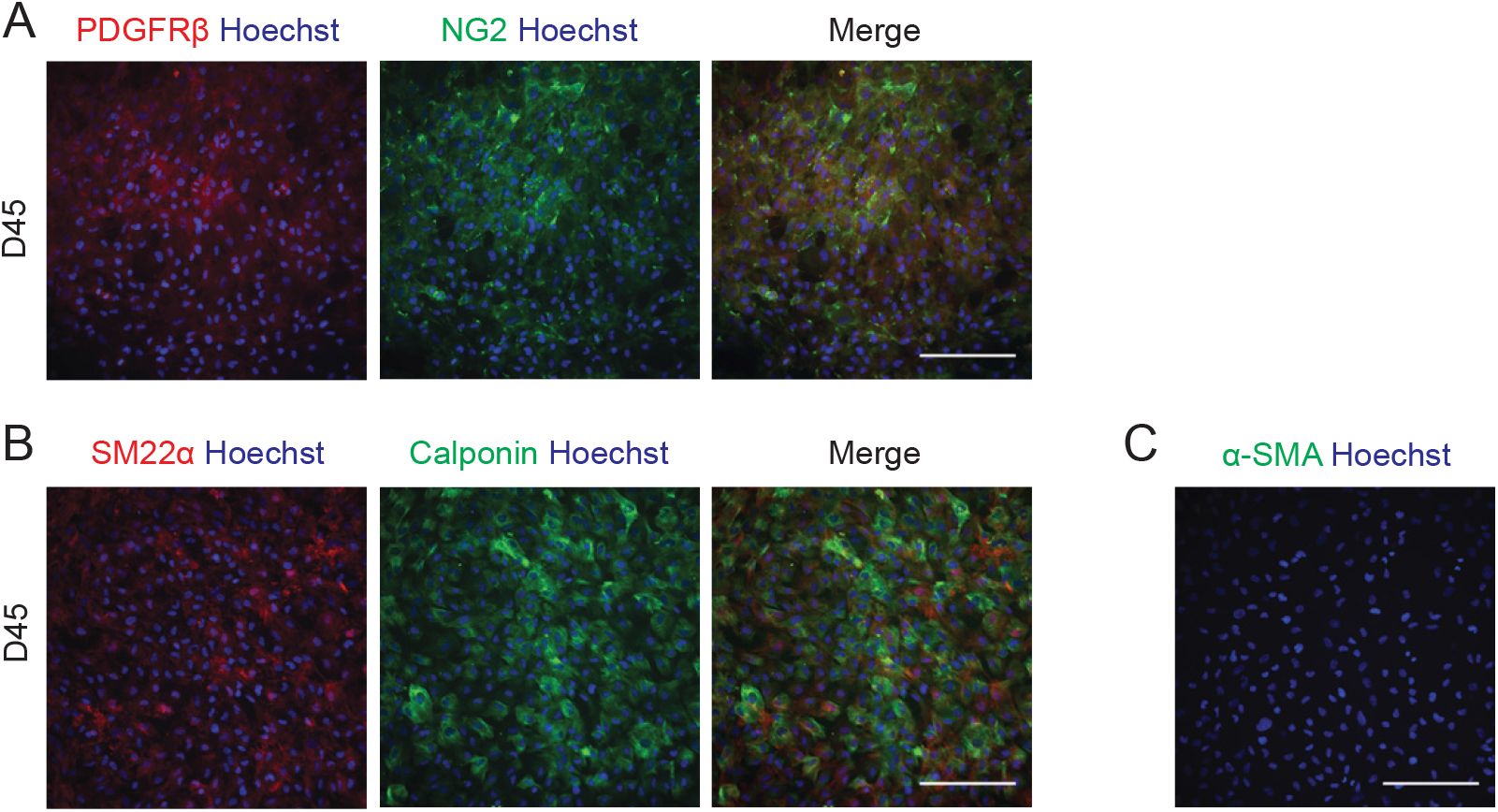
Long-term maintenance of hPSC-derived pericyte-like cells. A) PDGFRβ and NG2 immunocytochemistry of H9-derived pericyte-like cells maintained in E6 + 10% FBS to D45. Hoechst nuclear counter stain (blue) is also included. Scale bar: 200 μm. B) Calponin and SM22α immunocytochemistry of H9-derived pericyte-like cells maintained in E6 + 10% FBS to D45. Hoechst nuclear counter stain (blue) is also included. Scale bar: 200 μm. C) α-SMA immunocytochemistry of H9-derived pericyte-like cells maintained in E6 + 10% FBS to D45. Hoechst nuclear counter stain (blue) is also included. Scale bar: 200 μm.

**Figure S6:**
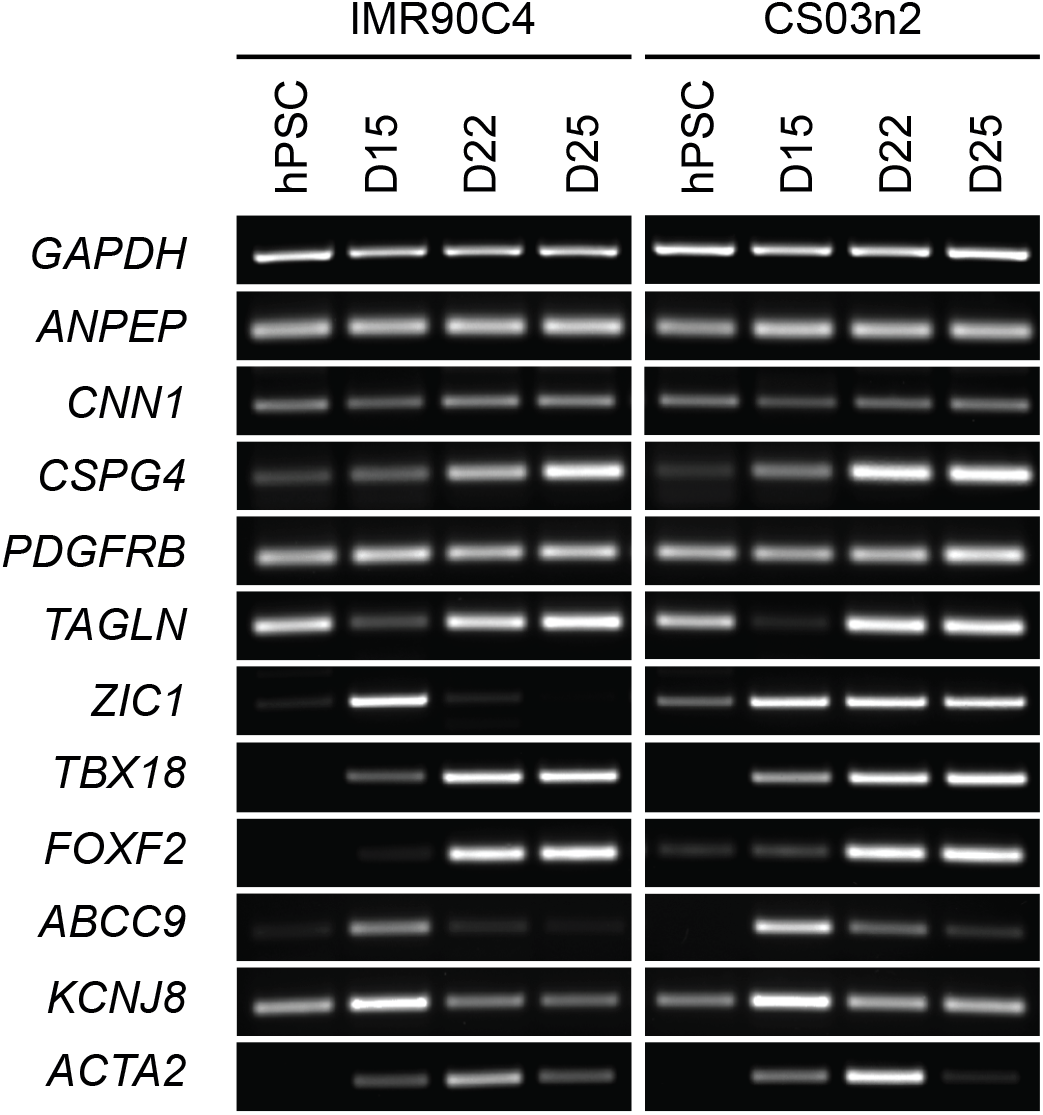
Temporal PCR analysis of mural and pericyte transcripts for the differentiating IMR90C4 and CS03n2 iPSC lines.

**Figure S7:**
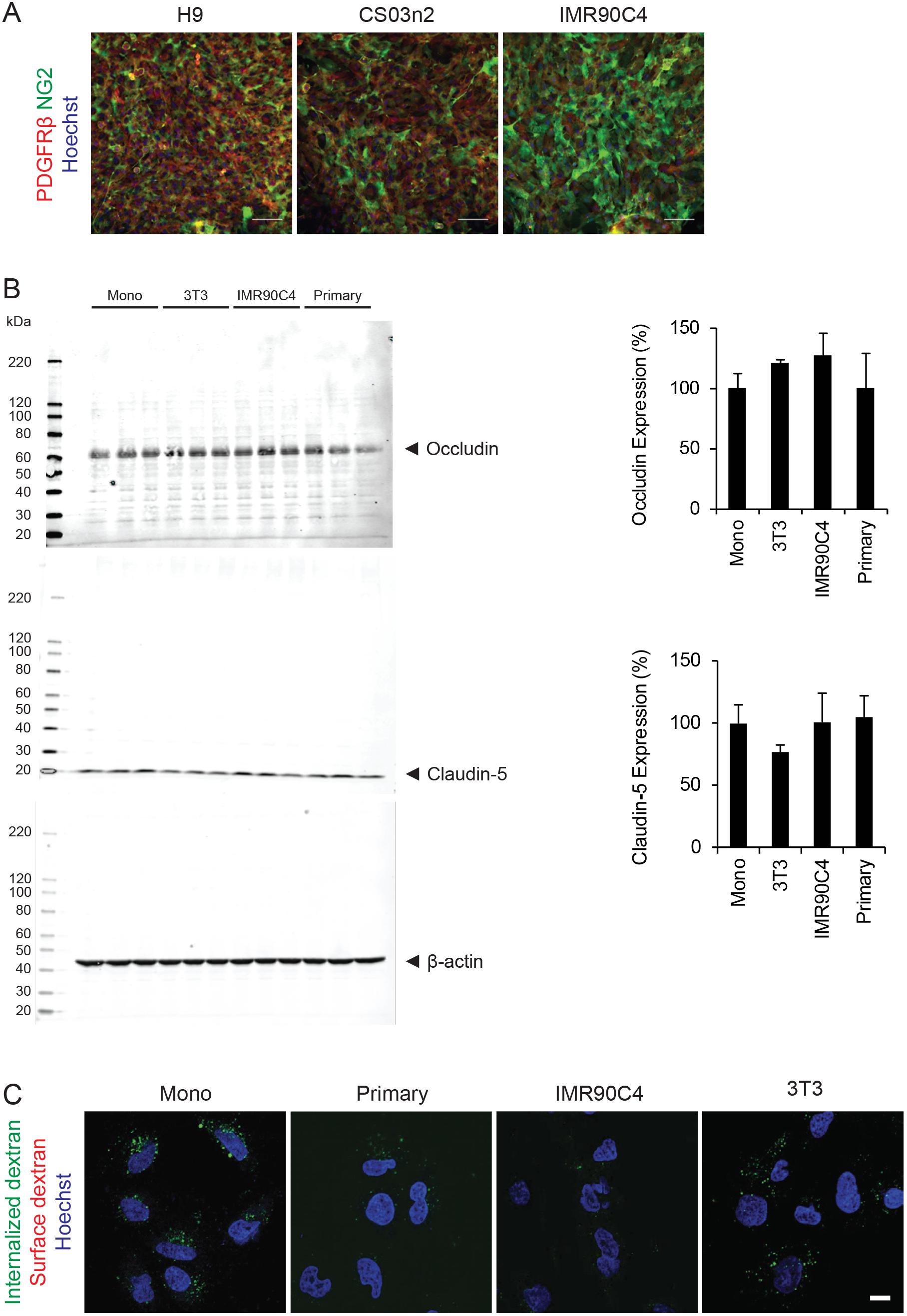
Supplemental analysis of BMEC/hPSC-derived pericyte-like cell co-cultures. A) PDGFRβ and NG2 immunocytochemistry of hPSC-derived pericyte-like cells following 48 hours of co-culture with iPSC-derived BMECs. Hoechst nuclear counter stain (blue) is also included. Images are representative of two independent differentiations. Scale bars: 100 μm. B) Western blot analysis of occludin and claudin-5 expression in iPSC-derived BMECs cultured alone or co-cultured with primary brain pericytes, IMR90C4-derived pericyte-like cells, or 3T3s. Quantification of occludin and claudin-5 expression after normalized to β-actin signal and to monoculture expression levels. Plotted are the means ± SD from 3 Transwells from a single differentiation. No significant differences by ANOVA. C) Confocal microscopy of monocultured iPSC-derived BMECs incubated with Alexa 488-tagged 10 kDa dextran (green) with EC medium (Mono) or conditioned medium from primary brain pericytes, IMR90C4-derived pericyte-like cells, or 3T3s. Total dextran is depicted in green. Surface dextran was labeled with Alexa 647 (red), with little observed signal. Thus, the observed green signal is a result of internalized dextran. Hoechst nuclear counter stain (blue) is also included. Scale bar: 10 μm.

**Figure S8:**
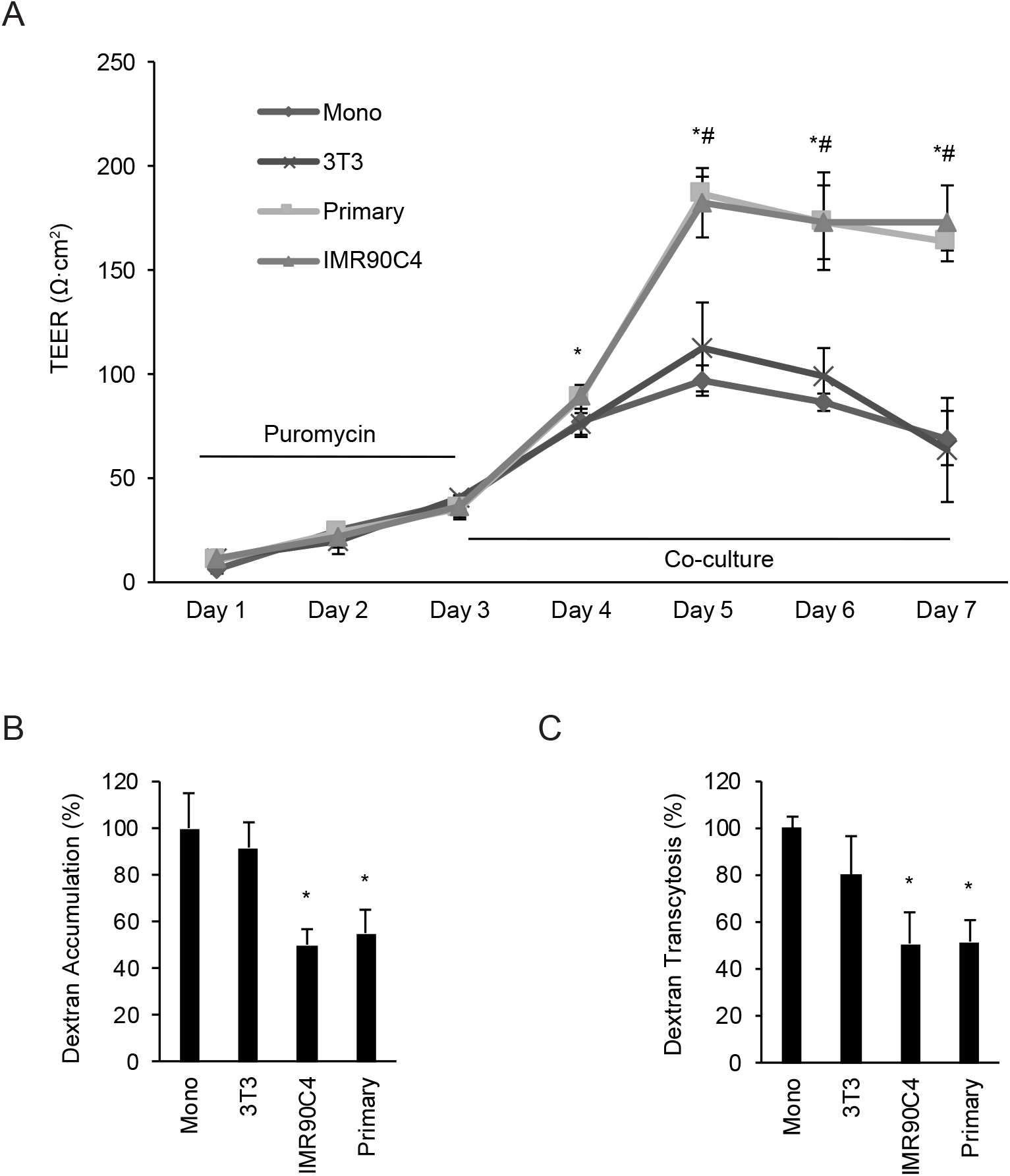
Measurement of the effects of hPSC-derived pericyte-like cells on primary rat BMEC phenotypes. A) TEER profile of primary rat BMECs either in monoculture or co-culture with primary brain pericytes, IMR90C4-derived pericyte-like cells, or 3T3s. Plotted are means ± SD of three Transwells from a single rat BMEC isolation. * *P* < 0.05 IMR90C4-derived pericyte-like cell co-culture vs. monoculture; # *P* < 0.05 primary pericyte co-culture vs. monoculture; ANOVA followed by Dunnett’s test. B,C) Accumulation (B) or transcytosis (C) of Alexa 488-tagged 10 kDa dextran in primary rat BMECs following co-culture with cell types as described in A. All results normalized to BMEC monoculture control. Plotted are the means ± SD of 3 Transwells from a single rat BMEC isolation. * *P* <0.05 vs. monoculture; ANOVA followed by Dunnett’s test.

**Table S1:**
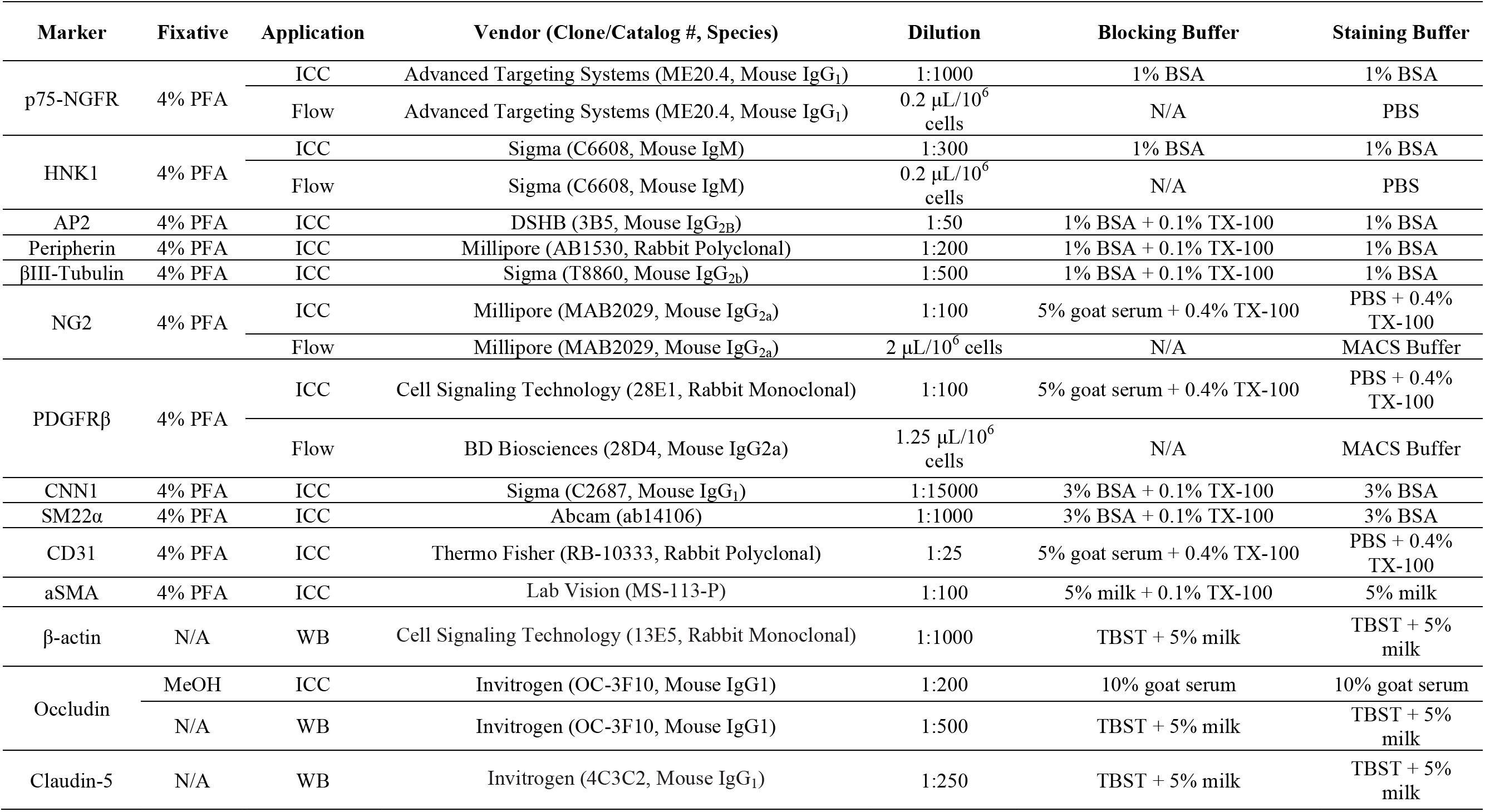
Antibody Staining Conditions.

| Marker | Fixative | Application | Vendor (Clone/Catalog #, Species) | Dilution | Blocking Buffer | Staining Buffer |
| --- | --- | --- | --- | --- | --- | --- |
| p75-NGFR | 4% PFA | ICC | Advanced Targeting Systems (ME20.4, Mouse IgG<sub>1</sub>) | 1:1000 | 1% BSA | 1% BSA |
| | | Flow | Advanced Targeting Systems (ME20.4, Mouse IgG<sub>1</sub>) | 0.2 μL/10<sup>6</sup> cells | N/A | PBS |
| HNK1 | 4% PFA | ICC | Sigma (C6608, Mouse IgM) | 1:300 | 1% BSA | 1% BSA |
| | | Flow | Sigma (C6608, Mouse IgM) | 0.2 μL/10<sup>6</sup> cells | N/A | PBS |
| AP2 | 4% PFA | ICC | DSHB (3B5, Mouse IgG<sub>2B</sub>) | 1:50 | 1% BSA + 0.1% TX-100 | 1% BSA |
| Peripherin | 4% PFA | ICC | Millipore (AB1530, Rabbit Polyclonal) | 1:200 | 1% BSA + 0.1% TX-100 | 1% BSA |
| βIII-Tubulin | 4% PFA | ICC | Sigma (T8860, Mouse IgG<sub>2b</sub>) | 1:500 | 1% BSA + 0.1% TX-100 | 1% BSA |
| NG2 | 4% PFA | ICC | Millipore (MAB2029, Mouse IgG<sub>2a</sub>) | 1:100 | 5% goat serum + 0.4% TX-100 | PBS + 0.4% TX-100 |
| | | Flow | Millipore (MAB2029, Mouse IgG<sub>2a</sub>) | 2 μL/10<sup>6</sup> cells | N/A | MACS Buffer |
| PDGFRβ | 4% PFA | ICC | Cell Signaling Technology (28E1, Rabbit Monoclonal) | 1:100 | 5% goat serum + 0.4% TX-100 | PBS + 0.4% TX-100 |
| | | Flow | BD Biosciences (28D4, Mouse IgG<sub>2a</sub>) | 1.25 μL/10<sup>6</sup> cells | N/A | MACS Buffer |
| CNN1 | 4% PFA | ICC | Sigma (C2687, Mouse IgG<sub>1</sub>) | 1:15000 | 3% BSA + 0.1% TX-100 | 3% BSA |
| SM22α | 4% PFA | ICC | Abcam (ab14106) | 1:1000 | 3% BSA + 0.1% TX-100 | 3% BSA |
| CD31 | 4% PFA | ICC | Thermo Fisher (RB-10333, Rabbit Polyclonal) | 1:25 | 5% goat serum + 0.4% TX-100 | PBS + 0.4% TX-100 |
| aSMA | 4% PFA | ICC | Lab Vision (MS-113-P) | 1:100 | 5% milk + 0.1% TX-100 | 5% milk |
| β-actin | N/A | WB | Cell Signaling Technology (13E5, Rabbit Monoclonal) | 1:1000 | TBST + 5% milk | TBST + 5% milk |
| Occludin | MeOH | ICC | Invitrogen (OC-3F10, Mouse IgG<sub>1</sub>) | 1:200 | 10% goat serum | 10% goat serum |
|  | N/A | WB | Invitrogen (OC-3F10, Mouse IgG<sub>1</sub>) | 1:500 | TBST + 5% milk | TBST + 5% milk |
| Claudin-5 | N/A | WB | Invitrogen (4C3C2, Mouse IgG<sub>1</sub>) | 1:250 | TBST + 5% milk | TBST + 5% milk |

**Table S2:**
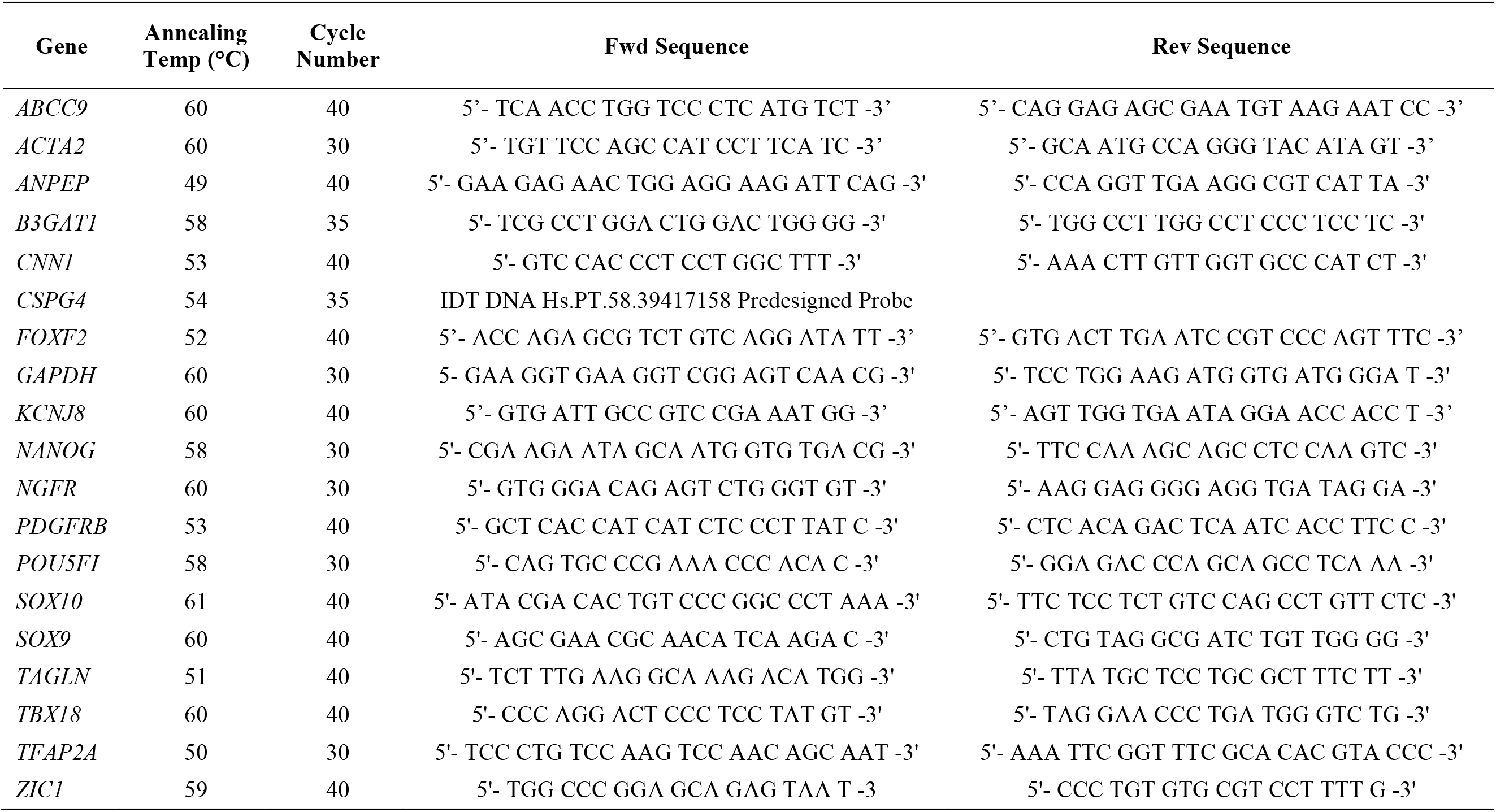
DNA Probe Sequences and Running Conditions.

**Table S3:** Pericyte-enriched genes identified by single cell RNA-sequencing in mouse (*39*) with human homologs.

| Gene | FPKM |  |
| --- | --- | --- |
|  | Primary | hPSC-derived* |
| *ABCC9* | 0.3 | 0.1 |
| *AGAP2* | 0.0 | 0.0 |
| *ANK2* | 9.6 | 2.3 |
| *ANO4* | 0.2 | 2.5 |
| *APOD* | 0.2 | 0.6 |
| *APOE* | 42.9 | 25.6 |
| *ARHGAP31* | 3.4 | 7.6 |
| *CORO1B* | 68.4 | 56.1 |
| *ECE1* | 58.7 | 31.0 |
| *EMCN* | 3.3 | 1.7 |
| *FAM118B* | 0.0 | 0.0 |
| *FBLN1* | 24.7 | 112.0 |
| *FLT1* | 4.7 | 7.6 |
| *FOXF2* | 4.4 | 0.9 |
| *GGT1* | 104.8 | 181.9 |
| *GPR4* | 5.4 | 4.6 |
| *IFI30* | 18.4 | 17.2 |
| *IGF2* | 0.3 | 14.0 |
| *ITIH5* | 2.6 | 0.2 |
| *ITM2A* | 0.3 | 3.9 |
| *JUP* | 6.4 | 20.8 |
| *KCNJ8* | 0.8 | 0.3 |
| *LGALS9* | 5.1 | 3.4 |
| *NODAL* | 0.1 | 0.1 |
| *NRXN2* | 2.5 | 0.8 |
| *NXPH4* | 30.3 | 0.4 |
| *PCDHGC3* | 0.0 | 0.0 |
| *PDE2A* | 0.6 | 0.1 |
| *PDE8A* | 6.5 | 6.0 |
| *PHC1* | 0.0 | 10.8 |
| *PHLDB1* | 28.5 | 28.8 |
| *PLOD1* | 189.1 | 145.8 |
| *POR* | 33.4 | 28.4 |
| *PPP1CC* | 40.7 | 82.8 |
| *PREX2* | 0.0 | 0.0 |
| *SEPP1 (SELENOP)* | 19.8 | 1.4 |
| *SFRP2* | 0.0 | 16.7 |
| *SLC22A8* | 0.0 | 0.0 |
| *SLC6A13* | 0.4 | 0.1 |
| *SPP1* | 0.96 | 11.6 |
| *ST8SIA4* | 0.0 | 0.3 |
| *SYNE1.2 (SYNE1)* | 16.9 | 6.0 |
| *TNFRSF19* | 0.4 | 13.6 |
| *TRPC3* | 0.2 | 0.1 |
| *TTLL3* | 160.3 | 210.9 |
| *UCHL1* | 739.1 | 415.9 |
| **Number with FPKM $\geq 1$** | **26** | **29** |

\* Average of all D25 hPSC-derived pericyte-like cell differentiations (H9 A-C, CS03n2 and IMR90C4)

## Abbreviations

BBB: blood-brain barrier
BMECs: brain microvascular endothelial cells
CNS: central nervous system
E6-CSFD: E6 medium supplemented with CHIR99021, SB431542, FGF2, and dorsomorphin
EGM-2: endothelial growth factor medium 2
hESCs: human embryonic stem cells
hPSCs: human pluripotent stem cells
iPSCs: induced pluripotent stem cells
NCSC: neural crest stem cells
NVU: neurovascular unit
vSMCs: vascular smooth muscle cells

